# Dynamic pneumococcal genetic adaptations support bacterial growth and inflammation during coinfection with influenza

**DOI:** 10.1101/659557

**Authors:** Amanda P. Smith, Lindey C. Lane, Tim van Opijnen, Stacie Woolard, Robert Carter, Amy Iverson, Corinna Burnham, Peter Vogel, Dana Roeber, Gabrielle Hochu, Michael D.L. Johnson, Jonathan A. McCullers, Jason Rosch, Amber M. Smith

## Abstract

*Streptococcus pneumoniae* (pneumococcus) is one of the primary bacterial pathogens that complicates influenza virus infections. These bacterial coinfections increase influenza-associated morbidity and mortality through a number of immunological and viral-mediated mechanisms, but the specific bacterial genes that contribute to post-influenza pathogenicity are not known. Here, we used genome-wide transposon mutagenesis (Tn-Seq) to reveal bacterial genes that confer improved fitness in influenza-infected hosts. The majority of the 32 identified genes are involved in bacterial metabolism, including nucleotide biosynthesis, amino acid biosynthesis, protein translation, and membrane transport. We generated single-gene deletion (SGD) mutants of five identified genes: SPD1414, SPD2047 (*cbiO1),* SPD0058 (*purD*), SPD1098, and SPD0822 (*proB*), to investigate their effect on *in vivo* fitness, disease severity, and host immune responses. Growth of SGD mutants was slightly attenuated *in vitro* and *in vivo*, but each still grew to high titers in the lungs of mock- and influenza-infected hosts. Despite high bacterial loads, mortality was significantly reduced or delayed with all SGD mutants. Time-dependent reductions in pulmonary neutrophils, inflammatory macrophages, and select proinflammatory cytokines and chemokines were also observed. Immunohistochemical staining further revealed that neutrophil phenotype and distribution was altered in the lungs of influenza-SGD coinfected animals. These studies demonstrate a critical role for specific bacterial genes and for bacterial metabolism in driving virulence and modulating immune function during influenza-associated bacterial pneumonia.

## Introduction

Bacterial pathogens often complicate influenza virus infections, causing increased morbidity and mortality, and *Streptococcus pneumoniae* (pneumococcus) is one of the leading pathogens that has presented as a risk factor for hospitalization, severe disease, and mortality during infuenza epidemics and pandemics (1–7). There is considerable genetic diversity within and between pneumococcal serotypes, and laboratory and clinical studies suggest that specific strains are preferentially promoted in influenza-infected hosts (8–10). Despite the clinical importance of the synergy between influenza viruses and pneumococci, a systematic investigation into bacterial factors that contribute to coinfection disease severity has yet to be performed.

During influenza A virus (IAV) coinfection with pneumococcus, bacteria are able to grow rapidly, viral burden increases, and significant inflammation amasses. The host-pathogen interplay is complex with numerous factors contributing to pathogen growth and host disease. Several studies have investigated the impact of viral virulence factors (e.g., reduced epithelial integrity, PB1-F2, viral neuraminidase (NA)), bacterial virulence factors (e.g., bacterial NA, enhanced attachment, sialic acid catabolism), and host immune responses (e.g., cytokine storm, cell dysfunction and death) on the pathogenicity of bacterial coinfections during influenza virus infection (11–17). Incidence and severity of coinfection is, in part, a function of the detrimental effects that influenza virus infection has on key immune responses (e.g., alveolar macrophages (AMΦ) depletion and dysfunction (18–23), neutrophil and inflammatory macrophage dysfunction (23–32), and abundant production of pro-inflammatory cytokines (12–17, 33)). However, the role that single genes play in pathogenicity during coinfection remains unclear.

Pneumococci, in particular, are highly adaptable and alter gene expression and metabolic functions to adjust to multiple host niches during infection and several genes have been identified as regulators of transmissibility and disease severity during primary pneumococcal infection (34–45). Select pneumococcal virulence factors have been investigated in the context of IAV infection (e.g., sialic acid catabolism) (46–49), but systematic genomic screens have not yet been employed to assess pneumococcal adaptations during IAV-pneumococcal coinfection. Genome-wide screens have been used to identify influenza-induced changes during *Haemophilus influenzae* coinfection (50), which included changes in purine biosynthesis, amino acid metabolism, iron homeostasis, and cell wall synthesis (50). Genomic screens assessing pneumococcal adaptations of the TIGR4 strain in hosts with sickle cell anemia suggested genes involved in purine biosynthesis, complement function, and iron acquisition were of importance (51). Given these findings and the similarities in bacterial metabolic adaptations under various host pressures, understanding how bacterial genes influence influenza-pneumococcal coinfection is imperative.

Here, we used genome-wide transposon insertion sequencing (Tn-Seq) (52) to investigate all non-essential pneumococcal genes that modulate disease severity and immune responses. Doing so could help establish important species- and strain-dependent and independent mechanisms amenable to targeting with therapeutics. Our Tn-Seq screen revealed 32 total genes, some with known metabolic functions, that confer bacterial fitness during influenza-pneumococcal coinfection compared to primary bacterial infection. Interestingly, there were time-dependent differences. To determine how select genes affect pathogenicity and host immune responses, we generated 5 single-gene deletion (SGD) mutants (D39Δ*cbiO1* (SPD2047), D39Δ*purD* (SPD0058), D39Δ*1414*, D39Δ*1098*, and D39Δ*proB* (SPD0822)). Lethality was significantly reduced or eliminated during infection in infected animals despite high bacterial loads in the lungs and blood. This was concurrent with significant reductions in innate pulmonary immune responses and development of pneumonia in IAV-infected animals. Taken together, these data indicate a critical role for pneumococcal metabolism in shaping host responses and altering disease severity during post-influenza bacterial pneumonia.

## Results

### Bacterial Adaptations During Pneumococcal Coinfection With Influenza

To identify the pneumococcal factors that increase bacterial fitness during IAV infection, we employed a high-throughput, transposon sequencing (Tn-Seq) approach (52). We generated a library of ~50,000 transposon insertion mutants in the type 2 pneumococcal strain D39, then infected groups of mice with 75 TCID_50_ (50% tissue culture infectious dose) influenza A/Puerto Rico/34/8 (PR8) or mock control (PBS) followed 7 d later with 10^6^ CFU (colony forming units) of the transposon library. Bacteria were collected from lungs harvested at 12 h or 24 h post-bacterial infection (pbi), genomic DNA was isolated and sequenced, and the relative abundance and fitness of transposon mutants was calculated (Supplementary Information). In comparing the results from pre- and post-infection and from mock- and IAV-infected animals, 17 genes conferred differential fitness at 12 h, and 23 genes had differential fitness at 24 h pbi. Of these, 8 genes were detected at both time points (Table 1, Fig 1).

**Table 1:** Pneumococcal Genes with Significant Differential Fitness During Coinfection with IAV. Pneumococcal fitness genes identified at 12 h and 24 h pbi by Tn-Seq during IAV infection compared to mock-infected hosts. Highlighted in bold are the genes chosen for additional characterization.

| Locus | Gene | Description | 12 | 24 |
| --- | --- | --- | --- | --- |
| SPD_0002 | dnaN | DNA polymerase III subunit beta | √ |  |
| SPD_0052 |  | phosphoribosylformylglycinamide synthase, putative |  | √ |
| SPD_0053 | purF | amidophosphoribosyltransferase |  | √ |
| SPD_0054 | purM | phosphoribosylformylglycinamide cyclo-ligase |  | √ |
| SPD_0055 | purN | phosphoribosylglycinamide formyltransferase | √ | √ |
| SPD_0057 | purH | bifunctional purine biosynthesis protein PurH | √ | √ |
| **SPD_0058** | **purD** | **phosphoribosylamine--glycine ligase** | √ | √ |
| SPD_0059 | purE | phosphoribosylaminoimidazole carboxylase, catalytic | √ | √ |
| SPD_0182 |  | conserved hypothetical protein | √ |  |
| SPD_0372 |  | sodium:alanine symporter | √ | √ |
| SPD_0395 | efp | translation elongation factor P |  | √ |
| SPD_0723 | rpiA | ribose 5-phosphate isomerase A | √ |  |
| **SPD_0822** | **proB** | **glutamate 5-kinase** |  | √ |
| SPD_0823 | proA | gamma-glutamyl phosphate reductase |  | √ |
| SPD_0824 | proC | pyrroline-5-carboxylate reductase |  | √ |
| SPD_0907 | hemK | HemK protein | √ |  |
| SPD_0994 | ribF | riboflavin biosynthesis protein RibF | √ |  |
| SPD_1087 | fhs | formate--tetrahydrofolate ligase |  | √ |
| SPD_1090 |  | membrane protein, putative |  | √ |
| **SPD_1098** |  | **amino acid ABC transporter, amino acid-binding** | √ | √ |
| SPD_1099 |  | amino acid ABC transporter, ATP-binding protein |  | √ |
| SPD_1130 | licD2 | phosphotransferase LicD2 |  | √ |
| SPD_1293 |  | acetyltransferase, GNAT family protein |  | √ |
| SPD_1309 | pgdA | peptidoglycan GlcNAc deacetylase |  | √ |
| SPD_1333 |  | conserved hypothetical protein |  | √ |
| SPD_1354 |  | conserved hypothetical protein | √ |  |
| **SPD_1414** |  | **oxalate:formate antiporter** | √ | √ |
| SPD_1468 |  | phosphoglycerate mutase | √ |  |
| SPD_1782 | ksgA | dimethyladenosine transferase | √ |  |
| SPD_1923 |  | 2,3,4,5-tetrahydropyridine-2-carboxylate N- | √ |  |
| **SPD_2047** | **cbiO1** | **cobalt ABC transporter, ATP-binding protein** | √ | √ |
| SPD_2048 | cbiO2 | cobalt ABC transporter, ATP-binding protein CbiO2 |  | √ |
| **Total** |  |  | **17** | **23** |

**Fig 1:**
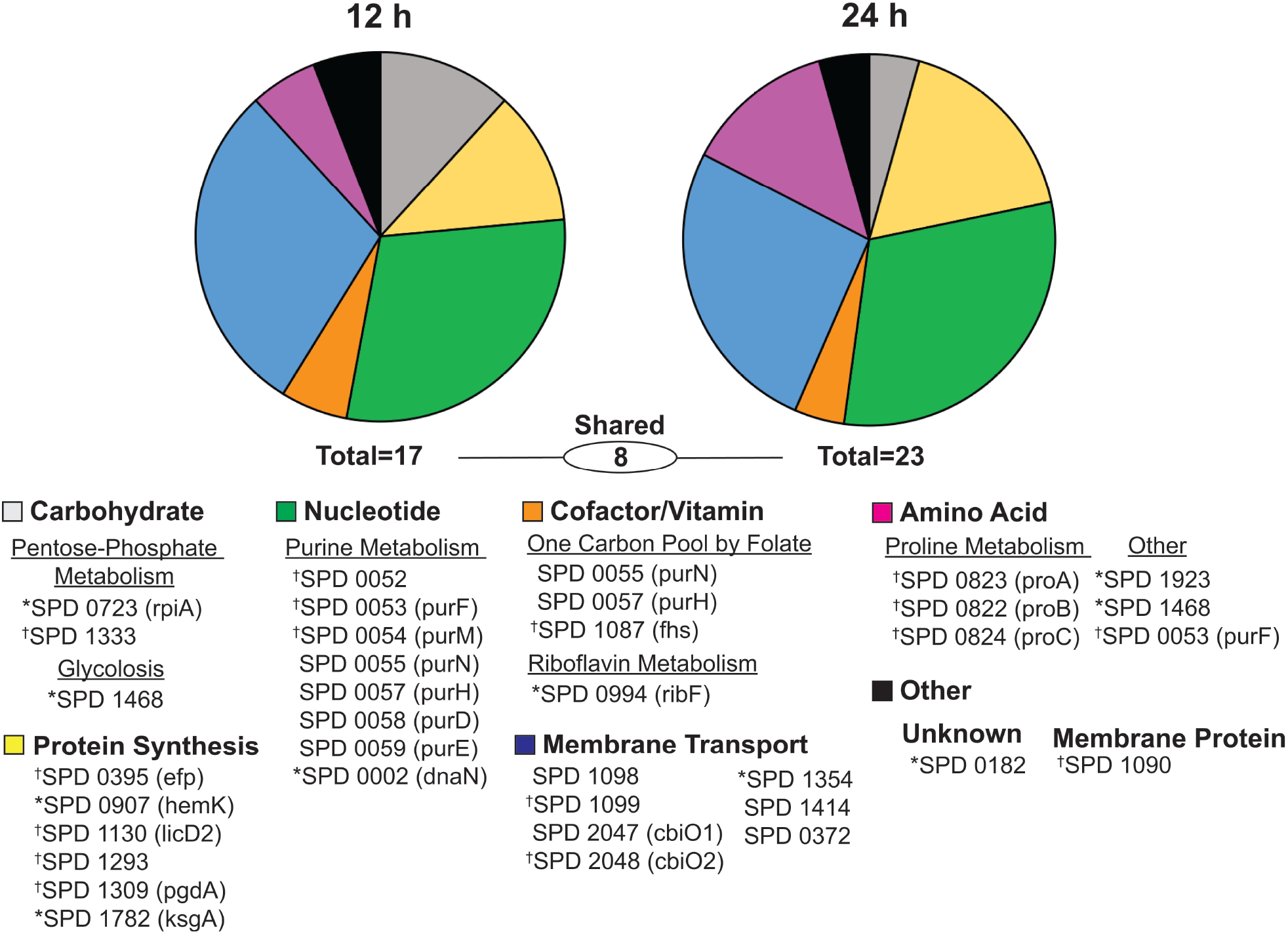
Time-Dependent Analysis of Pneumococcal Genes Impacting Fitness During Influenza Virus Coinfection. Functional breakdown of the pneumococcal genes that impact bacterial fitness during coinfection with influenza A virus as identified by Tn-Seq. Markers identify genes important at only *12 h pbi (17 total) or only at ^†^24 h pbi (23 total). No marker indicates significance at both time points (8 total).

The core set of genes identified at 12 h and 24 h pbi are responsible for amino acid biosynthesis, nucleotide biosynthesis, protein translation, and membrane transport (Fig 1). Genes in the purine biosynthesis pathway comprised the largest number of genes (8 total; SPD0002 (*dnaN*), SPD0052, SPD0053 (*purF*), SPD0054 (*purM*), SPD0055 (*purN*), SPD0057 (*purH*), SPD0058 (*purD*), and SPD0059 (*purE*)) followed by ATP-binding cassette (ABC) transporters (5 total; SPD1098, SPD1099, SPD2047 (*cbiO1)*, SPD2048 (*cbiO2*), and SPD1354 (putative)), protein translation (6 total; SPD0395 (*efp*), SPD1782 (*ksgA*), SPD0907 (hemK), SPD1130 (licD2), SPD1293, SPD1923), and proline biosynthesis (3 total; SPD0822 (*proB*), SPD0823 (*proA*), SPD0824 (*proC*)). Other genes included a putative membrane protein (SPD1090), carbon metabolism (SPD0723 (*ripA*), SPD1087, SPD1333 (putative), and SPD1468), and riboflavin metabolism (SPD0994).

### Impaired Bacterial Metabolism Selectively Reduces Fitness *In Vitro*

We generated single-gene deletion (SGD) bacterial mutants from 3 of the identified categories (D39Δ*cbiO1*, D39Δ*1414*, D39Δ*1098* (membrane transport), D39Δ*purD* (purine metabolism), and D39Δ*proB* (proline metabolism)) to assess the differential fitness of genes predicted by Tn-seq (Table 1, Fig 1, and Table S1). Growth of the SGD mutants in culture medium was unaffected, with the exception of D39Δ*cbiO1*, which was significantly attenuated from 3 h to 8 h (p<0.05) and was the only SGD strain that had not autolysed after 24 h (Fig 2A). During metabolic starvation (i.e., PBS), D39Δ*1098* decayed more rapidly (p=0.01) and D39Δ*cbiO1* more slowly (p<0.01), but other SGD mutants were similar to WT (Fig 2B). Bacterial growth was rescued when PBS cultures were supplemented with lung homogenate supernatants (s/n) (Fig S1A-B) and was not different than WT D39 after 6 h of culture in lung homogenate s/n from mock- or IAV-infected mice (75 TCID_50_ PR8, 7 d) (Fig S1C-D).

**Fig 2:**
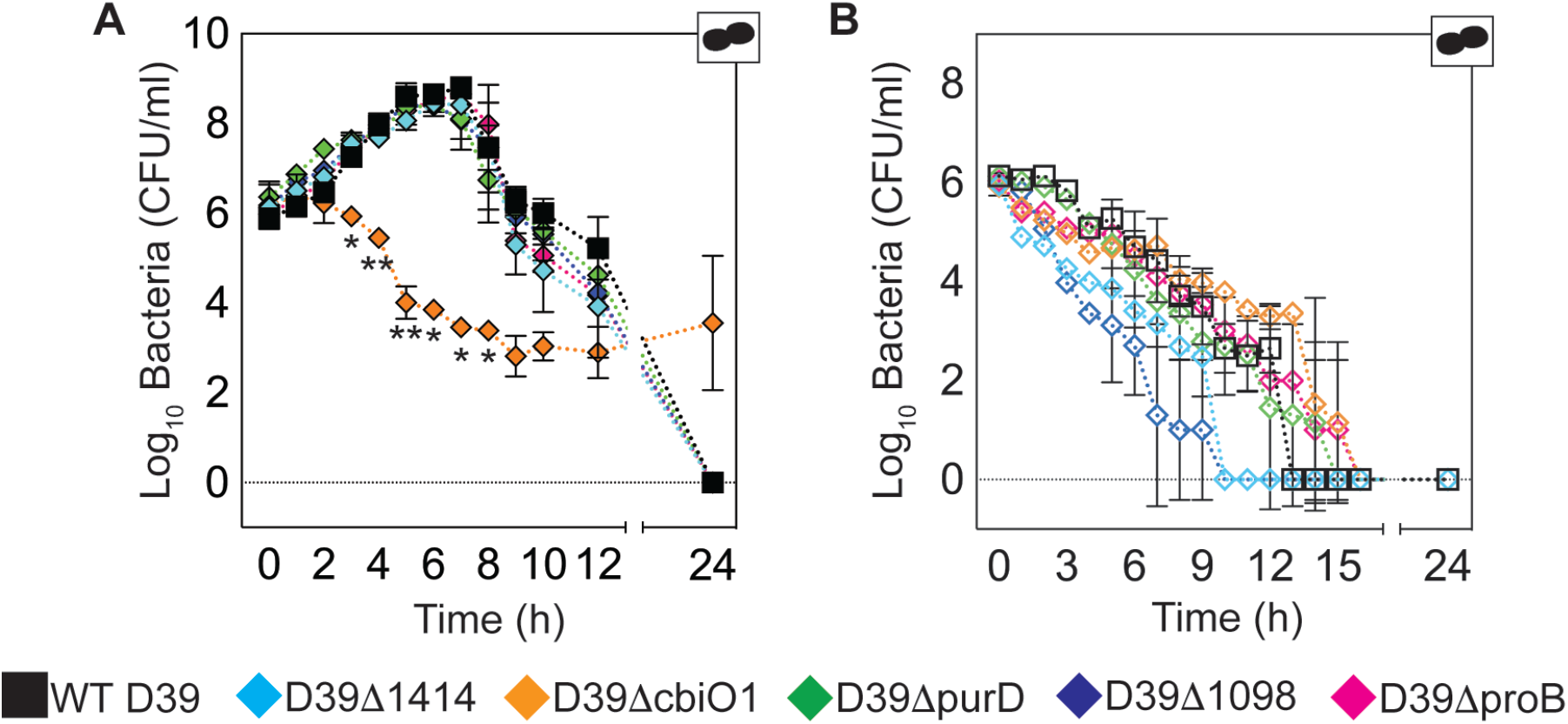
*In vitro* Growth Kinetics of SGD Mutant Bacteria. Bacteria were grown at 37°C in 1 ml of ThyB (A) or PBS (B). Signficance is indicated as *p<0.05 and **p<0.01 (D39Δ*cbiO1* only) compared to WT D39 at the indicated time point (Panel A). Each symbol (squares or diamonds) is the mean of two representative experiment and the bars are the geometric mean ± standard deviation (SD). Cartoons indicate pathogens in the culture (bacteria alone).

### Impaired Bacterial Metabolism Protects Against Virulence *In vivo* Reduced Mortality

To establish the effect of gene deletion on pathogenicity, weight loss was monitored as a measure of disease severity (Fig 3, Table S2). In mock- and IAV-infected animals, infection with WT D39 resulted in 100% mortality by 72 h pbi and 48 pbi, respectively (Fig 3A-B). In mock-infected animals, mortality was reduced by 90-100% in 4 out of 5 SGD mutants and by 40% with D39Δ*proB* (all p<0.01) (Fig 3A, Table S2). Correspondingly, weight loss was significantly reduced (48 h pbi; p<0.05) (Fig 3C). In IAV-infected animals, mortality was reduced by 40-90% in 4 out of 5 SGD mutants (all p<0.01) (Fig 3B, Table S2). Coinfection with D39Δ*proB* resulted in 100% mortality; however, the mean survival time was lengthened by 5 d (p<0.01) (Fig 3B, Table S2). In IAV-infected animals, weight loss was not significantly reduced compared to WT D39 (24 h pbi; p>0.05) (Fig 3D) and weights of all SGD-infected animals, except D39Δ*proB*, began to rebound ~3 d pbi and returned to baseline by 12-14 d pbi.

**Fig 3:**
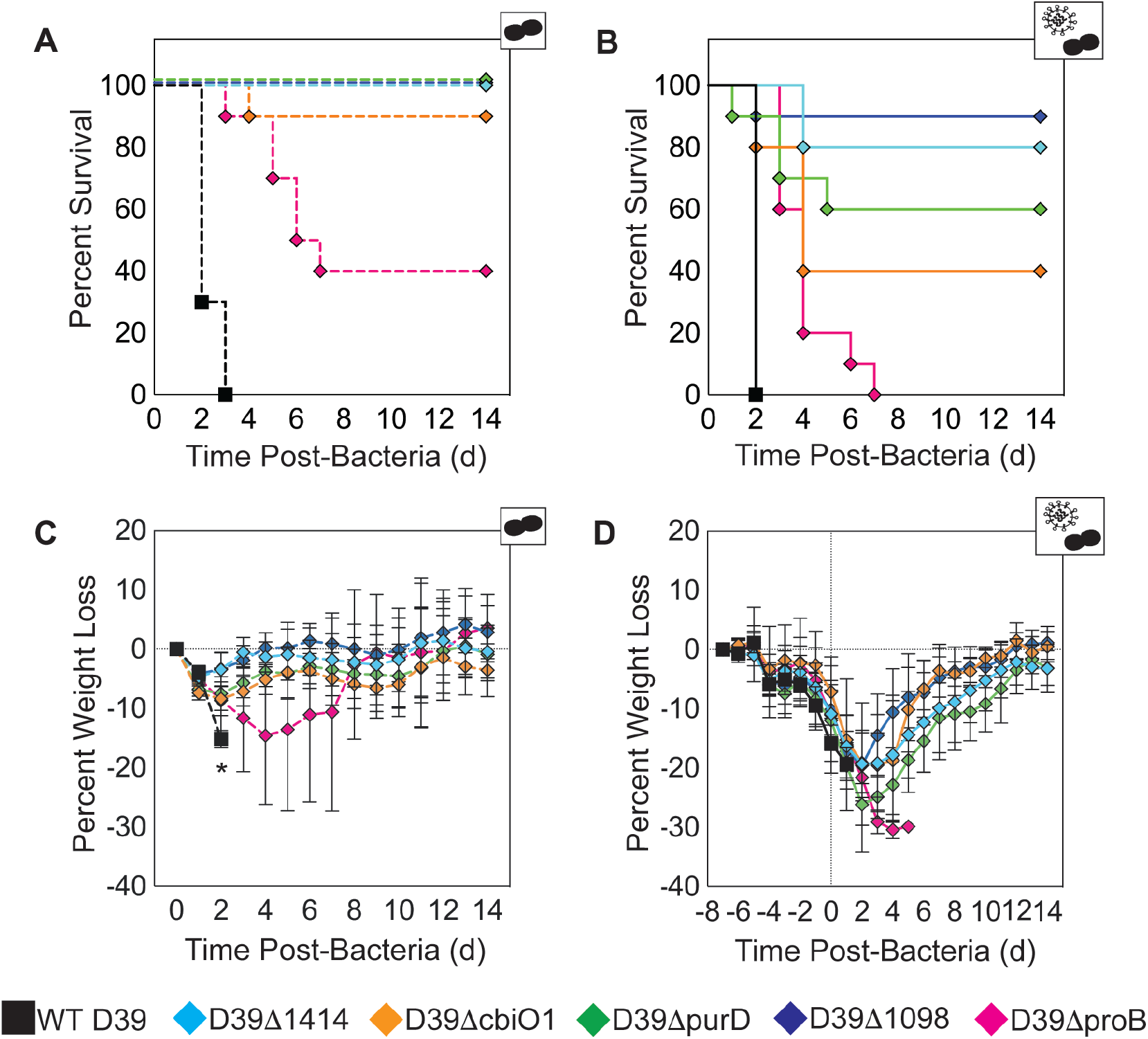
*In vivo* Pathogenicity of Infection with SGD Mutant Bacteria. Kaplan-Meier survival curves (A-B) and weight loss (percent loss compared to naïve) (C-D) of mice mock-infected (PBS) (dashed lines; Panel A, C) or IAV-infected (75 TCID_50_ PR8) (solid lines; Panel B, D) followed 7 d later with 10^6^ CFU of the indicated bacteria. Survival curves are significantly different (p<0.01) for each SGD mutant compared to WT D39, in mock- (Panel A) and IAV- (Panel B) infected animals. Differences in survival curves are detailed in Table S2. Signficant difference in weight loss is indicated as *p<0.05 for each SGD mutant (diamonds) compared to WT D39 (square) at the indicated time point (Panel C). Data are shown as the mean ± standard deviation (SD) from 10 mice/group. Cartoons indicate infection status of study group (bacteria alone or virus plus bacteria).

### Reduced Bacteria in the Lung and Blood

Lung titer kinetics of the SGD mutants *in vivo* (Fig 4A-B) mirrored *in vitro* titer kinetics in lung s/n supplemented cultures (Fig S2C-D). At 4 h pbi, SGD mutant lung titers were 0.6-1.4 log_10_ CFU lower than WT D39 in mock-infected animals and 0.4-2.1 log_10_ CFU lower in IAV-infected animals (p<0.01) (Fig 4A-B, Table S2). By 24 h pbi, titers were statistically similar, yet reduced compared to WT D39 in the lungs (0.5-2.7 log_10_ CFU reduction; p>0.05) and blood (1.3-6.6 log_10_ CFU reduction; p>0.05) of mock-infected animals (Fig 4A, Fig 4C, and Table S2). However, in IAV-infected animals, SGD mutant titers remained significantly lower in the lungs (0.8-2.1 log_10_ CFU reduction; p<0.05) and blood (by 2.2-5.0 log_10_ CFU reduction; p<0.01) at 24 h pbi (Fig 4B, Fig 4D, and Table S2). Despite attenuated growth in the lungs and reduced bacteremia at 24 h pbi, bacterial loads for each SGD mutant remained high in the lungs and blood of coinfected animals at 24 h, 48 h, and 72 h pbi (Fig 4B, Fig 4D, and Table S2).

**Fig 4:**
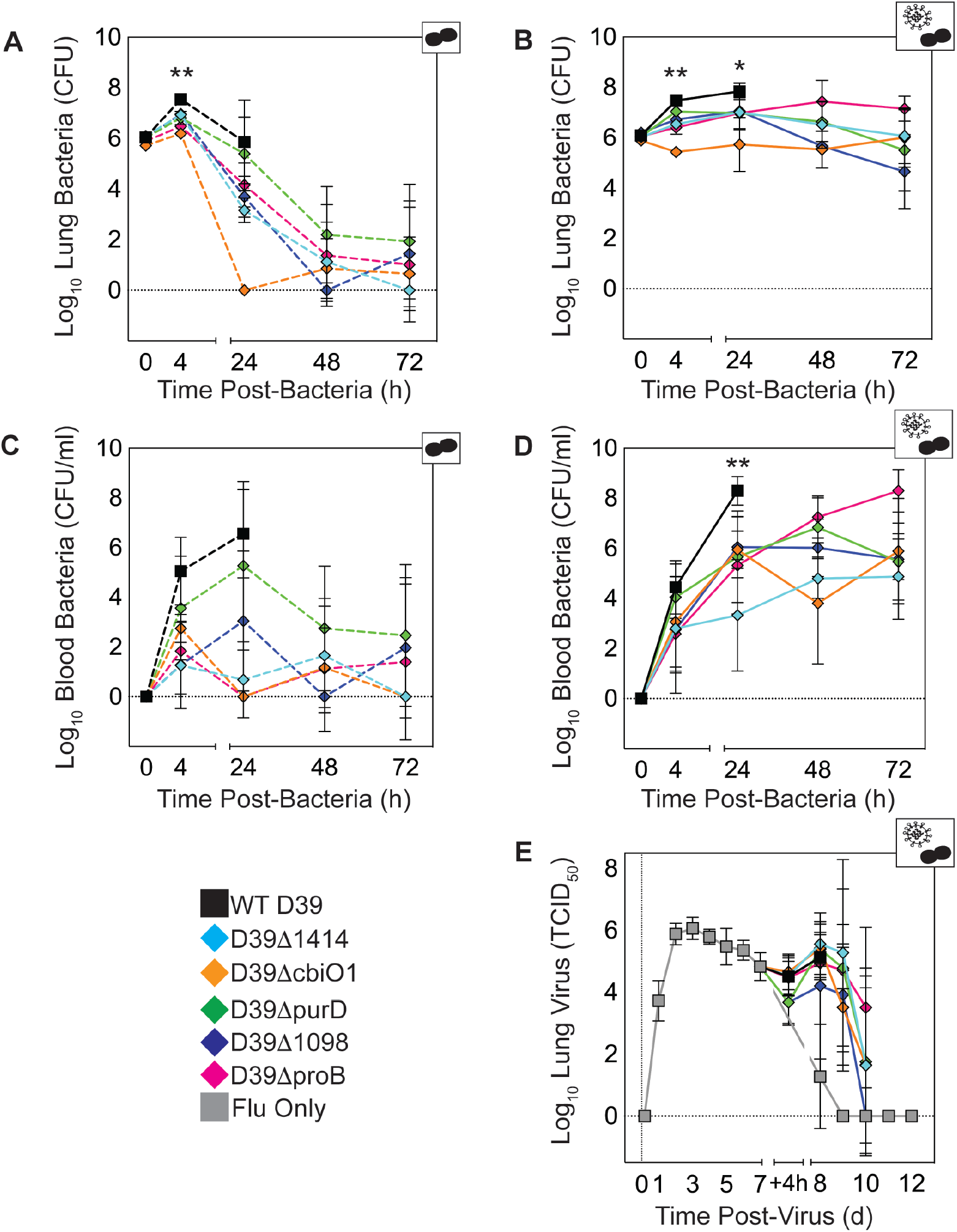
Viral and Bacterial Titer Kinetics. Lung bacterial titers (A-B), blood bacterial titers (C-D), and lung viral titers (E) from groups of mice either mock-infected (PBS) (dashed lines; Panels A, C) or IAV-infected (75 TCID_50_ PR8) (solid lines; Panels B, D, E) followed 7 d later with 10^6^ CFU of the indicated bacteria. Signficance is indicated as *p<0.05 and **p<0.01 for each of the SGD mutants compared to WT D39 at the indicated time point. Data are shown as the mean ± standard deviation (SD) from 5 mice/group. Cartoons indicate infection status of study group (bacteria alone or virus plus bacteria). The log_10_ change in pathogen loads between each SGD mutant and WT D39, and analysis of variance results are summarized in Table S2.

### Similar Viral Load Kinetics

Similar to previous studies (53), viral loads rebounded following coinfection with each SGD mutant bacteria. While viral rebound and clearance occurred with varying dynamics, each was statistically similar to coinfection with WT D39 (p>0.05) (Fig 4E, Table S2).

### Altered Cytokine and Chemokine Responses

To better understand the reduced morbidity and mortality during infection with the SGD mutants, we examined cytokine and chemokine dynamics in the lungs (Fig 5, Fig S2-S4). In IAV-infected animals, IFN-α, IFN-β, IL-6, KC, MIP-1β and GM-CSF were significantly reduced at 24 h pbi with SGD mutant bacteria compared to WT D39 (p<0.05), with the exception of IFN-α during coinfection with D39Δ*1414* (p=0.95) and D39Δ*purD* (p=0.17) (Fig 5, Fig S2-S4). Interestingly, at 4 h pbi in IAV-infected animals, IFN-α was elevated in D39Δ*1414*, D39Δ*cbiO1*, D39Δ*purD*, and D39Δ*1098* coinfections (p<0.01) and IFN-β was elevated in D39Δ*cbiO1* coinfection (p<0.01) (Fig 5, Fig S2J and L). In mock-infected animals, IFN-α, IL-6, KC, and MIP-1β were similar at 24 h pbi (p>0.05) (Fig 5, Fig S2-S4). IL-1α, IL-1β, MIP-1α, and TNF-α were reduced in both mock- and IAV-infected animals at 24 h pbi with each SGD mutant compared to WT D39 (p<0.05), except for D39Δ*purD* in mock-infected animals (p>0.05) (Fig 5, Fig S2-S4). Minimal differences were detected for MCP-1, RANTES, IL-2, IFN-γ, IL-10, IL-12(p40), and IL-12(p70), except during primary infection with D39Δ*1098* and D39Δ*proB* (Fig 5, Fig S3-S4).

**Fig 5:**
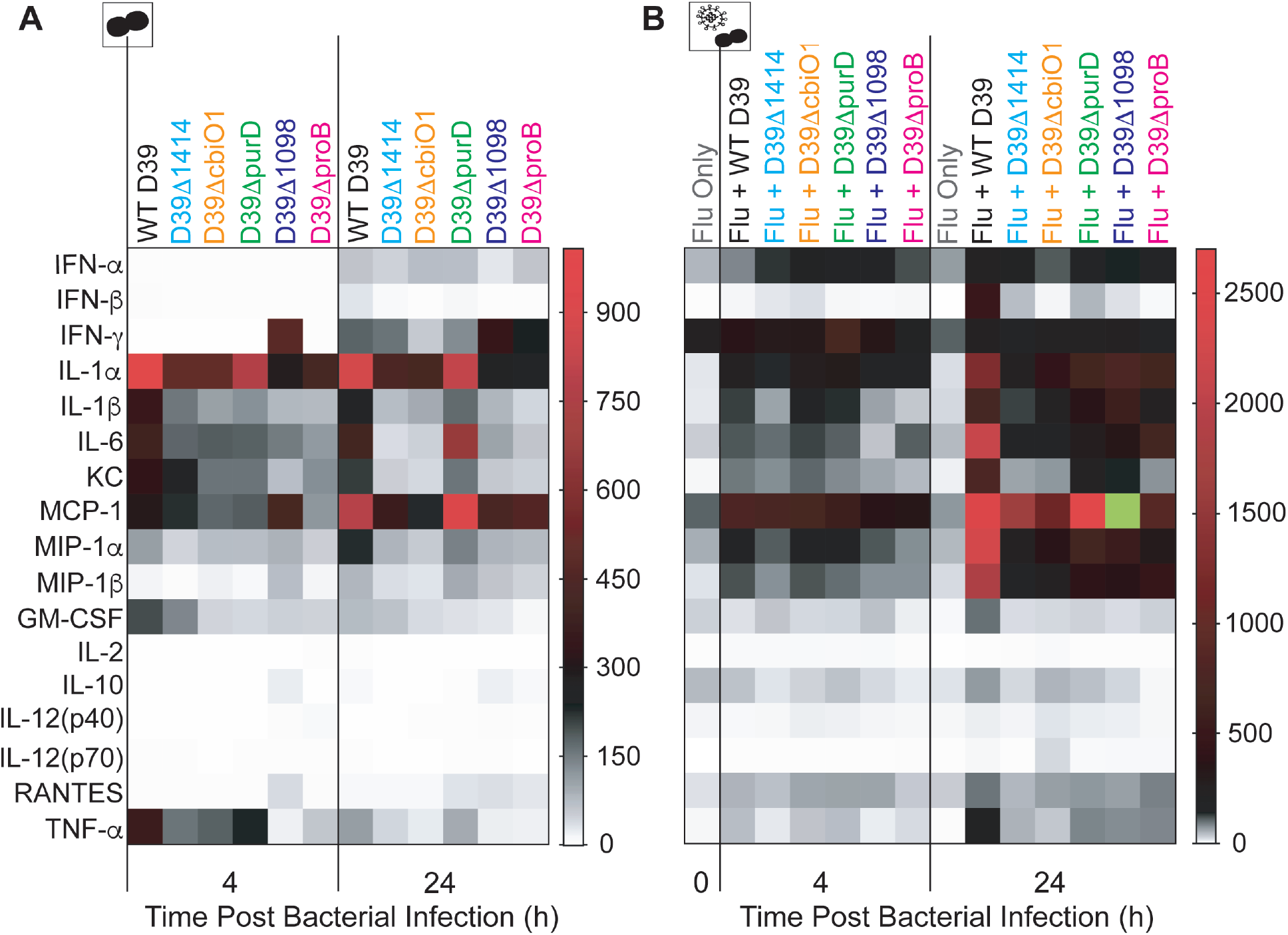
Heat Map of Cytokine and Chemokine Changes. Fold change compared to naïve in the mean value of the indicated cytokine/chemokine at 4 h or 24 h pbi in the lungs of mice mock-infected (PBS) (A) or IAV-infected (75 TCID_50_ PR8) (B) followed 7 d later with 10^6^ CFU of the indicated bacteria. The green square is ouside of the listed range and indicates a 4021 fold change. Cartoons indicate infection status of study group (bacteria alone or virus plus bacteria). Plots depicting absolute log_10_ picograms (pg) of measured cytokines and chemokines are in Figs S2-S4.

### Reduced Inflamatory Cell Responses

In accordance with the changes detected in pulmonary cytokines and chemokines, infection with SGD mutant bacteria altered the dynamics of select immune cells in the lungs (Fig 6 and Figs S5-S6). In IAV-infected animals, neutrophils (Ly6G^hi^) were significantly reduced at 4 h pbi with D39Δ*1098* (p<0.01) and D39Δ*proB* (p<0.05), and at 24 h pbi with D39Δ*1414*, D39Δ*cbiO1*, D39Δ*purD*, and D39Δ*proB* (p<0.05) (Fig 6B). Inflammatory macrophages (IMΦs; Ly6G^−^, CD11c^hi^, F4/80^hi^, CD11b^+^) were also reduced at 4 h pbi during coinfection with D39Δ*1414*, D39Δ*purD*, D39Δ*1098*, and D39Δ*proB* (p<0.05) (Fig 6D). D39Δ*cbiO1* did not lead to reduced IMΦs at 4 h pbi (p=0.14), but did induce a significant increase in IMΦs at 24 h pbi (p<0.01) (Fig 6D). In mock-infected animals, neutrophils and IMΦs were not significantly different at 4 h pbi for any of the SGD mutants (p>0.05) (Fig 6A, Fig 6C). At 24 h pbi, neutrophils were reduced by D39Δ*purD* (p<0.05) and IMΦs were increased by D39Δ*cbiO1* (p<0.05) infection (Fig 6A, Fig 6C). There were minimal differences in T cell populations (Fig S6I-L) or the extent of AMΦ depletion (Ly6G^−^, CD11c^hi^, F4/80^hi^, CD11b^−^) (Fig 6E-F) in mock- or IAV-infected animals.

**Fig 6:**
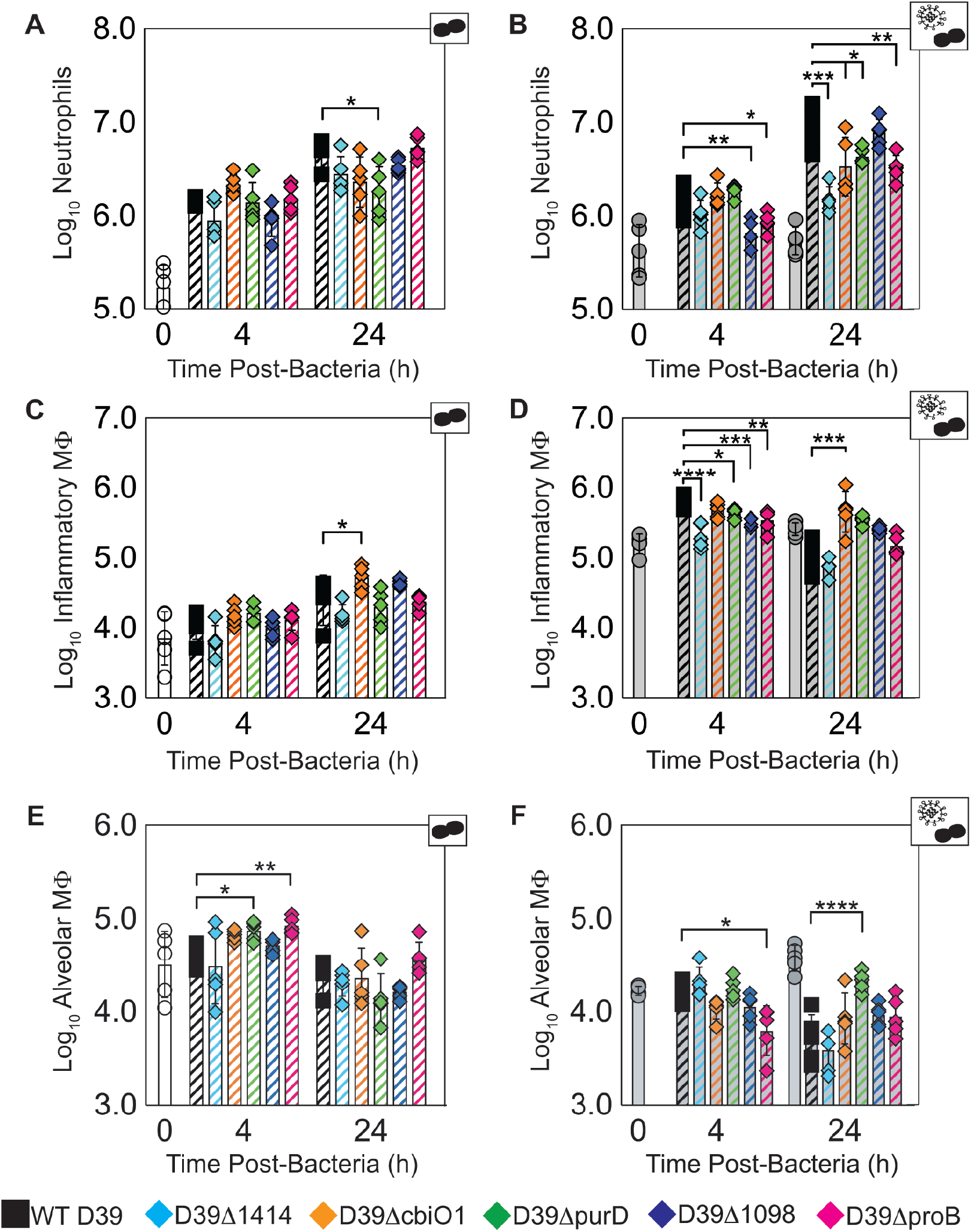
Pulmonary Immune Cell Kinetics. Kinetics at 4 h or 24 h pbi of neutrophils (A-B), inflammatory macrophages (C-D), and alveolar macrophages (E-F) from mice mock-infected (PBS) (Panels A, C, E) or IAV-infected (75 TCID_50_ PR8) (Panels B, D, F) followed 7 d later with 10^6^ CFU of the indicated bacteria. Each symbol (circles, squares, or diamonds) represents a single mouse, and the bars are the geometric mean ± standard deviation (SD)) from 5 mice/group. Mice were either uninfected (open white), IAV-infected only for 7 d (T=0) or 8 d (T=24) (solid grey), bacteria-infected (open hashed, colored), or IAV-bacteria coinfected (solid hashed, colored). Signficance is indicated as *p<0.05, **p<0.01, ***p<0.005, ****p<0.0001. Cartoons indicate infection status of study group (bacteria alone or virus plus bacteria). The flow cytometry gating scheme is in Fig S5 and additional cellular dynamics are in Fig S6.

### Reduced Pathology and Altered Neutrophil Phenotype

There was extensive pulmonary consolidation at 24 h pbi in the lungs of mice coinfected with WT D39, as characterized by thickened septa and alveoli filled with a mixture of neutrophils, free bacteria, and proteinaceous exudates (Fig 7A). These lesions were dramatically reduced during coinfection with D39Δ*proB*, D39Δ*1098*, and D39Δ1414 and almost absent in D39Δ*cbiO1* and D39Δ*purD* coinfection (Fig 7A). Immunohistochemical (IHC) staining showed a similar extent of influenza antigen (Fig 7B) but dramatically less bacterial antigen at 24 h pbi during coinfection with each of the SGD mutants (Fig 7C, Fig S7A), which mirrored the pulmonary viral and bacterial loads (Fig 4B, Fig 4E). In WT D39 coinfected animals, intracellular and extracellular bacterial antigen was present throughout influenza-lesioned areas, including pervascular connective tissues, consolidated alveolar parenchyma, and the central hypocellular area of resolving lesions. Bacterial antigen was not detected in the resolving influenza lesions with any of the SGD mutants, except for D39Δ*1098* where few bacteria or pneumococcal antigen-positive macrophages were present. IHC staining also showed massive neutrophil infiltration in WT D39 coinfection, and the resolving influenza lesions were consistently surrounded by sharply demarcated hypercellular bands composed of mixed viable and degenerating neutrophils (Fig 7C). In contrast, during D39Δ*proB*, D39Δ*purD*, and D39Δ*1098* coinfections, neutrophilic infiltrates generally appeared in clusters within alveoli peripheral to influenza lesions or scattered widely throughout inflamed areas in the lungs. Neutrophil infiltrates in these lungs consisted mostly of intact non-degenerating cells that formed indistinct bands surrounding the influenza lesions (Fig 7C). Animals coinfected with D39Δ*1414* and D39Δ*cbiO1* had the lowest number of neutrophils, which were mostly non-degenerating and scattered throughout inflamed areas (Fig 7C). Neutrophils were rare or absent within the resolving influenza lesions in animals coinfected with an SGD mutant.

**Fig 7:**
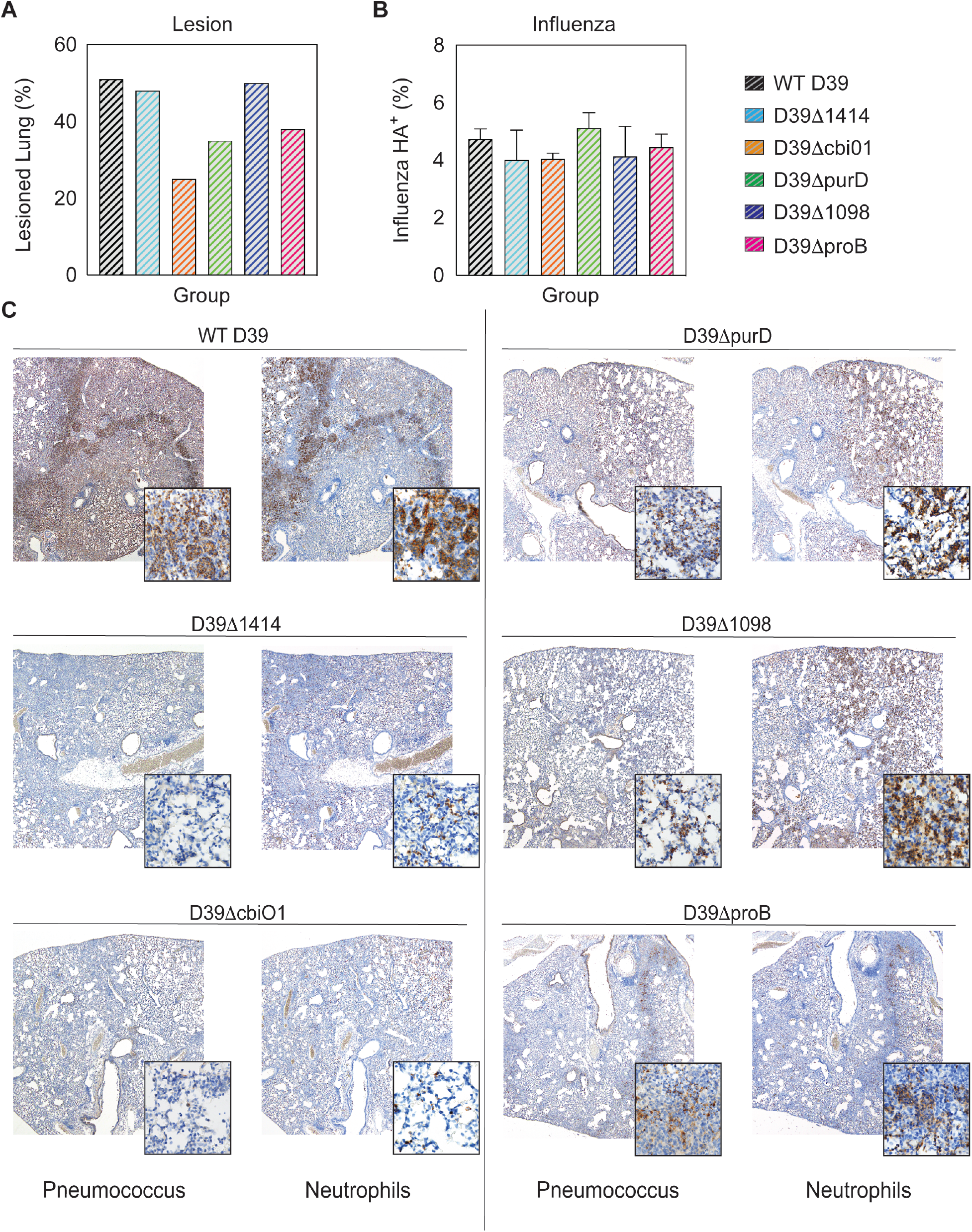
Lung Pathology During IAV-Pneumococcal Coinfection. Serial sections of lungs at 24 h pbi from mice IAV-infected (75 TCID_50_ PR8) followed 7 d later with 10^6^ CFU of the indicated bacteria, stained with hematoxylin and eosin (HE) for histological analysis (Panel A) or immunohistochemistry (IHC) for influenza HA glycoprotein (Panel B), pneumococcus, or neutrophil Ly6G/6C antigen (Panel C). Quantification was performed by a pathologist blinded to the study design (Panel A) or with MIPAR v3.0 image analysis software (Panel B and Figure S7). Representative images are at an original 4x magnification with 60x magnification insets.

## Discussion

Pathogenicity during IAV-pneumococcal coinfection is influenced by several pathogen and host factors, including the viral and bacterial strain and doses, and the strength of the inflammatory response (11–16). Our study here highlights the importance of specific genetic factors in contributing to this pathogenicity and to the dysregulated host response. By using Tn-Seq as an unbiased approach, we identified 32 genes as being critical to pneumococcal survival in the IAV-infected host in a time-dependant manner (Table 1, Fig 1). Unsurprisingly, known bacterial virulence genes were not varied in our screen, likely because those factors affect disease equally regardless of viral infection status (40, 41, 43). The identified genes are mostly involved in bacterial metabolism, which supports findings that the lung metabolome is altered during influenza virus infection. Our *in vivo* data suggest that some identified genes function as virulence factors in both mock- and IAV -infected hosts. However, the immunomodulatory effects differed dramatically between heathy mice and those infected with the influenza virus, highlighting the importance of immune and metabolic alterations in the susceptibility to and severity of bacterial coinfection.

Altered metabolic regulation has been observed during murine influenza virus infection (54–57), *in vitro* influenza virus infection of primary human bronchotracheal epithelial (HBAE) cells (58, 59), and in pediatric patients infected with influenza virus (58). However, even with the knowledge that influenza virus infection induces metabolic changes and that pneumococci modulate metabolic pathways under host-specific pressures (51, 60), the collection of genes identified here was not intuitive nor was their precise contribution to the infection dynamics. It has been shown that purine metabolic pathways are altered in *Haemophilus influenzae* and pneumococci within influenza-infected hosts (49, 50) and in pneumococci within hosts with sickle-cell disease (51). Here, eight genes with roles in purine metabolism were identified, including SPD0058 (*purD*) (61–63) (Table 1, Fig 1). This could indicate a common mechanism for bacterial adaptation within inflammatory environments. In influenza-infected hosts, purine biosynthesis is upregulated (55) and purine analogs can be used to reduce disease severity (e.g., by the antiviral T-705 (64, 65)). However, the intermediates of purine biosynthesis become depleted following bacterial infection (50), rendering genes of de novo purine intermediate synthesis (*purD* and *purC*) critical to coinfecting bacteria (50, 51) (Table 1, Fig 1). Here, it is likely that altered purine scavenging and competition for environmentally available purines/purine intermediates, modulates host immune cell function during infection with D39Δ*purD*, contributing to decreased lethality (Fig 3B) and immunopathology (Fig 7A) in IAV-infected hosts.

Several of the genes identified here act in glutamate/glutamine biosynthesis which, while important for bacterial pathogenesis in otherwise healthy individualts, plays a key role during influenza-pneumococcal coinfection. Here, both genes in the locus pinpointed as the main ABC glutamine/glutamate transporter of pneumococci (SPD1098/1099) (66, 67) were identified in our screen (Table 1, Fig 1). Deleting SPD1098 increased survival by 90% in coinfected animals (Fig 3B), despite neutrophil accumulation equivalent to that in WT D39 coinfection (Fig 6B, Fig 7C). Genes involved in proline biosynthesis, a process downstream of glutamate metabolism (61–63), were also identified, including SPD0822/0823 (*proB/A*) (Table 1, Fig 1). Deletion of SPD0822 (*proB*) delayed mortality during coinfection (Fig 3B) and led to reduced neutrophils and iMΦ in influenza-infected hosts, but not in mock-infected hosts (Fig 6). Glutamine is utilized at a high rate by immune cells and is needed for optimal function of macrophages and neutrophils (68–73). Since pulmonary cells have increased dependence on glutamine during influenza virus infection (57, 58), it is probable that competition for limited, enivonmentally available glutamine altered neutrophil infiltration and function, thereby reducing morbidity and mortality in IAV-infected hosts coinfected with these SGD mutants (Fig 5–7 and Fig 3B). Investigating metabolic alterations of innate immune cells during IAV-infection, would clarify the host-pathogen interactions observed here.

The functions of some genes identified here, including three other ABC transporters (e.g., SPD2047/2048 (*cbiO1/2* and SPD1414), are not well characterized. While it is understood that ABC transporters are important for pneumococcal virulence (35, 36, 74–77), investigating the impact of each gene identified in our screen provides insight into respiratory changes that occur during influenza virus infection. The role of cobalt in pneumococcal physiology and virulence remains unclear (78, 79), but the cbiO locus (putative cobalt transporter) was previously identified by microarray analysis as necessary for pneumococcal pathogenicity (42). Interestingly, bacteria lacking SPD2047 reach significantly higher lung titers in IAV-infected animals than in mock-infected animals (Fig 4A-D), suggesting that cobalt metabolism may be modified by influenza virus infection. To our knowledge, no study has assessed cobalt dynamics during influenza. The TIGR4 analog of the oxalate/formate antiporter SPD1414 (SP1587) has been shown to be important for bacterial survival in the lungs and blood (36, 80), cerebral spinal fluid (36), and nasopharynx (80). Here, deletion of SPD1414 reduced lethality (Fig 3B), neutrophil infiltration, and pulmonary damage in IAV-infected hosts (Fig 6–7). These results indicate roles for cobalt and oxalate/formate in the susceptibility to pneumococcal pneumonia during influenza-infection.

Mortality during influenza-pneumococcal coinfection is typically associated with an exuberant immune response coupled with high pathogen loads in the lung and the blood (14–16, 24, 81–83). However, reduced inflammation can lessen disease severity even with sustained bacterial loads (26). Here, growth of SGD mutant bacteria in IAV-infected animals was significantly attenuated (0.8-2.1 log_10_ reduction compared to WT D39), but not strictly correlated to pathogenicity and insufficient to account for the extreme reductions in mortality (up to 90%) (Fig 3B, Fig 4, Table S2) as equivalently reduced bacterial loads of WT D39 in IAV-infected animals still induce significant mortality (FigS8). In addition, AMΦ, which dictate the initial pneumococcal invasion and growth kinetics during IAV infection (18–20, 23), are not different during coinfection with most SGD mutants compared to WT D39 (Fig 6F). Contraction of the pulmonary CD8^+^ T cells (84) (Fig S6L) and the rebound of viral loads (Fig 4E) were not altered during coinfection with SGD mutant bacteria, which supports our previous findings that the mechanisms underlying rapid bacterial growth are independent from those that influence the post-bacterial viral rebound and pathogenicity (19, 20). These findings suggest that the observed improvements in immune responses and disease outcome are not solely driven by the differences in bacterial loads. Reduced cytokine levels, specifically lowered type I IFNs (Fig 5, Fig S2J and L) can improve neutrophil function (21, 24, 25, 85–89) and reduce epithelial cell death and lung permeability (90) and, thus, reduce pathogenicity (Fig 3B, Fig 5, and Fig 7). Moreover, reduced neutrophil infiltration and degeneration (Fig 6B, Fig 7C) may have mitigated the damaging cytokine storm that is typically associated with IAV-pneumococcal pneumonia (12–17, 33). A direct correlation between survival and any single host immune response in the early stages of coinfection (0-24 h pbi) was not readily apparent. It is possible that unmeasured components and/or cumulative effects influence lethality at later time points. These studies underscore the independent nature of pathogen growth and pathogenicity and illuminate the difficulty in reducing coinfection pathogenicity to a single variable.

Understanding how bacterial adaptations influence the development of pneumonia during influenza virus infections is important to effectively combat the disease. Here, we provide insight into the contribution of specific pneumococcal genes and to the regulatory host-pathogen dynamics that arise during IAV-pneumococcal coinfection. Our findings highlight the critical role of influenza-induced metabolic shifts in promoting bacterial infection and altering immune function, and suggest that targeting a single pneumococcal gene or metabolite could be an effective intervention to abrogate bacterial pneumonia during influenza infection. Further dissecting bacterial adaptation may identify additional therapeutic targets that could be used to prevent or treat post-influenza bacterial infections.

## Materials and Methods

### Ethics Statement

All experimental procedures were performed under protocol O2A-020 approved by the Animal Care and Use Committee at St. Jude Children's Research Hospital under relevant institutional and American Veterinary Medical Association (AVMA) guidelines and were performed in a biosafety level 2 facility that is accredited by the American Association for Laboratory Animal Science (AALAS).

### Mice

Adult (6 week old) female BALB/cJ mice were obtained from Jackson Laboratories (Bar Harbor, ME). Mice were housed in groups of 5 in high-temperature 31.2cm x 23.5cm x 15.2cm polycarbonate cages with isolator lids. Rooms used for housing animals were maintained on a 12:12-hour light:dark cycle at 22 ± 2°C with 50% humidity in the biosafety level 2 facility at St. Jude Children's Research Hospital (Memphis, TN). Prior to inclusion in the experiments, mice were allowed at least 7 days to acclimate to the animal facility such that they were 7 weeks old at the time of infection. Laboratory Autoclavable Rodent Diet (PMI Nutrition International, St. Louis, MO) and autoclaved water were available ad libitum. All experiments were performed under an approved protocol and in accordance with the guidelines set forth by the Animal Care and Use Committee at St. Jude Children’s Research Hospital.

### Tn-Seq

Plasmid DNA harboring *magellan6,* a derivative of the Himar1 Mariner transposon, was purified from *E. coli* with the Qiagen mini plasmid preparation kit (Qiagen). Pneumococcal DNA was isolated by phenol/chloroform extraction and ethanol precipitation from an exponentially growing culture in ThyB media (30 mg/ml Todd-Hewitt Broth powder and 0.2 mg/ml yeast). *In vitro, magellan6* transposition reactions were carried out with purified MarC9 transposase, 1 μg of pneumococcal target DNA and 1 μg of *magellan6* plasmid DNA. Reactions were incubated for 1 h at 30°C, inactivated for 20 min at 72°C, ethanol precipitated and resuspended in gap repair buffer [50 mM Tris (pH 7.8), 10 mM MgCl_2_, 1 mM DTT, 100 nM dNTPs and 50 ng BSA]. Repair of transposition product gaps was performed with *E. coli* DNA ligase overnight at 16°C. Repaired transposition products were transformed into naturally competent pneumococcal strain D39. The following day, colonies were scraped off tryptic soy-agar (TSA) plates supplemented with 3% sheep erythrocytes and 200 mg/ml spectinomycin (TSA-Spec), pooled into libraries of approximately 50,000 transformants/library, split up into multiple starter cultures and stored at −20°C.

### Bacterial Fitness By Tn-Seq

Bacteria were collected from the blood and lungs of each infected mouse at 12 h pbi or 24 h pbi and plated on TSA-Spec plates as indicated (Lung and Blood Harvesting). Later time points could not be examined due to insufficient numbers of surviving mice. Bacteria were incubated for 12 h at 37°C then collected in ThyB media and centrifuged at 4°C, 500xg for 10 min. The media supernatant was removed and the pellets were stored at −20°C. Gene identification by Tn-Seq was performed as described previously (51, 52, 80). The samples at each of the three time points (pre-selection (inoculum, t_1_) and post-selection (after infection, 12 h (t_2_) or 24 h (t_3_) pbi), were sequenced in rapid run mode on an Illumina HiSeq 2000 using the primers and cycle conditions in Table S1. For each insertion, the fitness *Wi*, was calculated by comparing the fold expansion of the mutant relative to the rest of the population with the following equation (91),

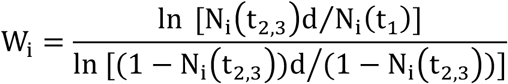

where the mutant frequency at time 0 and harvest are N_i_(t_1_) and N_i_(t_2,3_), respectively. The expansion factor (d) accounts for bacterial growth during library selection. Additional details of the method are included in the Supplementary Information.

### Infectious Agents

All experiments were done using the mouse adapted influenza A/Puerto Rico/8/34 (H1N1) (PR8) and type 2 pneumococcal strain D39 variants. Transposon mutant library used for infections was generated as described above (Tn-Seq). To generate the single-gene deletion (SGD) mutants, genomic DNA was isolated from WT D39 as above (Tn-Seq) and used as the template for gene SOEing PCR (92, 93). In brief, regions flanking the D39 locus targeted for deletion and containing overhangs complementary to the erythromycin (ERM) resistance cassette (*ermB*) (labeled regions A and B) were generated using the primer sets and cycle conditions indicated in Table S1B (EasyA PCR kit (Agilent Tech) and BioRad T100 thermal cycler). Products were purified using the Zymoclean gel DNA recovery kit (Zymo Research), recombined by PCR (TaKaRa PCR kit (Clontech Laboratories)) using the primers and cycle conditions in Table S1C, and transformed into D39. Transformed bacteria were selected after overnight growth on TSA plates containing 1 μg/ml ERM (TSA-ERM). Infection stocks were grown in ThyB media containing 1 μg/ml ERM, and stored at −80°C in 12% glycerol. ERM resistance cassette insertion and target locus deletion were confirmed by PCR with the primer sets in Table S1D.

### Infection Experiments

The viral infectious dose (TCID_50_) was determined by interpolation using the method of Reed and Muench (94) using serial dilutions of virus on Madin-Darby canine kidney (MDCK) cells. Bacterial infectious doses (CFU) were determined by serial dilutions on TSA (WT), TSA-ERM (SGD mutants), or TSA-Spec (Tn-seq) plates. Frozen stocks of inocula were diluted in sterile PBS and administered intranasally to groups of 5 (for kinetics) or 10 (for bacteria collection and survival) mice lightly anesthetized with 2.5% inhaled isoflurane (Baxter, Deerfield, IL) in a total volume of 100 μl (50 μl per nostril). Mice were inoculated with either PBS or 75 TCID_50_ PR8 at day 0 then with 10^6^ CFU of the transposon mutant library, D39, or indicated SGD mutant (in 100 μl), 7 days later. This dose of bacteria was chosen to ensure sufficient amounts of bacteria were recovered in Tn-Seq experiments and the CFU of each inoculum was confirmed as above. For bacterial sequencing, 500 μl of the inoculum was plated on TSA-Spec plates (100 μl/plate) (t_1_ in Bacterial Fitness By Tn-Seq). Animals were weighed at the onset of infection and each subsequent day to monitor illness and mortality. Mice were euthanized if they became moribund or lost 30% of their starting body weight.

### Lung and Blood Harvesting

Mice were euthanized by CO_2_ asphyxiation. For bacterial sequencing, lungs were perfused with 10 ml PBS, aseptically harvested, washed three times in PBS, and placed on ice in 500 μl PBS. The post-perfusion fluid (mixture of blood and PBS) was plated immediately on TSA-Spec plates (150 μl/plate). Lungs were then enzyme digested with collagenase (1 mg/ml, Sigma), and physically homogenized by syringe plunger against a 40μm cell strainer. Cell suspensions were centrifuged at 4°C, 500xg for 7 min and the supernatant was plated on TSA-spec plates (500 μl; 100 μl/plate). For *in vitro* SGD growth, lungs were homogenized (Omni TH-01 with 5mm flat blade) and centrifuged at 4°C, 500xg for 7 min. For *in vivo* kinetics, unperfused lungs were enzyme digested with collagenase as above and supernatants were used to quantify viral titers, bacterial titers, cytokine/chemokine levels (5 mice/group). Following red blood cell lysis, cells were washed in MACS buffer (PBS, 0.1 M EDTA, 0.01 M HEPES, 5 mM EDTA and 5% heat-inactivated FBS), counted with trypan blue exclusion using a Cell Countess System (Invitrogen, Grand Island, NY), and prepared for flow cytometric analysis as described below.

### *In vitro* Kinetics

Bacteria were grown at 37°C in 1.0 ml of ThyB media, PBS, or lung homogenate supernatants (s/n). At each time point, 50 μl was removed, serially diluted in PBS, and plated on TSA (WT) or TSA-ERM plates. Bacterial titers were normalized to the total volume. 3 biological replicates were performed.

### Lung and Blood Titers

For each mouse, viral titers were obtained using serial dilutions on MDCK monolayers, and bacterial titers were obtained using serial dilutions on TSA (WT) or TSA-ERM (SGD mutants) plates.

### Cytokines

Cytokines and chemokines were measured in lung supernatant by luminex (GM-CSF, IFN-γ, IL-1α, IL-1β, IL-2, IL-6, IL-10, IL-12(p40), IL-12(p70), KC, MCP-1, MIP-1α, MIP-1β, RANTES, and TNF-α) and ELISA (IFN-α,β). Prior to use, cell debris and aggregates were removed by centrifugation at 4°C, 400xg. Milliplex magnetic bead cytokine/chemokine plates (Millipore) were prepared according to manufacturer’s instructions. Analysis was done using a BioRad BioPlex (HTF System) and Luminex xPonent software. ELISAs for IFN-α and IFN-β (PBL Assay Science) were prepared according to the manufacturer’s instructions. Plates were read at 450 nm and analyzed using elisaanalysis.com. Mean concentrations of duplicate samples were calculated by construction of standard curves using a weighted 5PL and 4PL regression for the Milliplex and ELISA data, respectively. Absolute quantities of each cytokine/chemokine were calculated based on mean concentration of replicate samples normalized to the lung supernatant volume collected during tissue processing.

### Flow Cytometric Analysis

Flow cytometry (LSRII Fortessa; Becton Dickinson, San Jose, CA) was performed on single cell suspensions after incubation with 200 μl of 1:2 dilution of Fc block (human-γ globulin) on ice for 30 min, followed by surface marker staining with anti-mouse antibodies: CD11c (clone N418, eFluor450, eBioscience), CD11b (clone M1/70, Alexa700, BD Biosciences), Ly6G (clone 1A8, PerCp-Cy5.5, Biolegend), Ly6C (clone HK1.4, APC, eBioscience), F4/80 (clone BM8, PE, eBioscience), CD3e (clone 145-2C11, PE-Cy7, BD Biosciences or BV785, Biolegend), CD4 (clone RM4-5, PE-Cy5, BD Biosciences), CD8α (clone 53-6.7, BV605, BD Biosciences), CD49b (clone DX5, APC-Cy7, Biolegend or APC-e780, Affymetrix Inc) and MHC-II (clone M5/114.15.2, FITC, eBioscience). The data were analyzed using FlowJo 10.4.2 (Tree Star, Ashland, OR) where viable cells were gated from a forward scatter/side scatter plot and singlet inclusion. Following neutrophil exclusion (Ly6G^hi^), macrophages (MΦ) were gated as CD11c^hi^F4/80^hi^ with alveolar macrophages (AMΦ) sub-gated as CD11b^−^ and inflammatory macrophages (iMΦ) as CD11b^+^. After macrophage exclusion, T cell populations were gated as CD3e^+^ and subgated into CD8 T cells (CD3^+^CD8^+^CD4^−^DX5^−^) and CD4 T cells (CD3^+^CD8^−^CD4^+^DX5^−^). From the CD3e^−^ population, natural killer (NK) cells were gated as CD3^−^DX5^+^ and dendritic cells (DCs) as CD3^−^DX5^−^. DC were further gated into three subsets of DC; CD11c^+^CD11b^−^, CD11c^+^CD11b^+^, and CD11c^−^CD11b^+^ (Fig S5). The expression levels of MHC-II was used to confirm the identities and activation of MΦ and DC subsets. The absolute numbers of cell types were calculated based on viable events analyzed by flow cytometry and normalized to the total number of viable cells per sample.

### Histology

Mice were euthanized by CO_2_ asphyxiation and lungs were inflated *in situ* via tracheal infusion with 10% neutral-buffered formalin solution (NBF; ThermoFisher Scientific, Waltham, MA), followed by continued fixation in NBF for at least 2 weeks before being embedded in paraffin, sectioned at 4 μm, mounted on positively charged glass slides (Superfrost Plus; Thermo Fisher Scientific, Waltham, MA), and dried at 60°C for 20 min. Tissue sections were stained with hematoxylin and eosin (HE) or subjected to immunohistochemical (IHC) staining to detect influenza antigen, pneumococcus, and neutrophils. For detection of these targets, tissue sections underwent antigen retrieval in a prediluted Cell Conditioning Solution (CC1; Cat# 950-124; Ventana Medical Systems, Indianapolis, IN) for 32 min on a Discovery Ultra immunostainer (Ventana Medical Systems, Tucson, AZ). Primary antibodies included (i) a polyclonal goat antibody raised against the HA glycoprotein of B/Florida/04/2006 (Yamagata lineage) influenza virus diluted 1:2000 (Cat# I7650-05G, US Biologicals, Swampscott, MA), (ii) a rabbit polyclonal antibody to *Streptococcus pneumoniae* diluted 1:1000 (Cat# NB100-64502; Novus Biologicals, Littleton, CO), and (iii) a rat monoclonal antibody to neutrophils (Ly6G6C) diluted 1:50 (Cat# NB600-1387; Novus Biologicals, Littleton, CO). Binding was detected using OmniMap anti-Goat (#760-4647), anti-Rabbit (#760-4311), and anti-Rat (#760-4457) HRP (RUO) respectively (Ventana Medical Systems), with DISCOVERY ChromoMap DAB Kit (Ventana Medical Systems) as chromogenic substrate. Stained sections were examined by a pathologist (PV) blinded to the experimental group assignments. IHC was quantified (APS) using the provided immunohistochemistry recipe #L006-02 in MIPAR v3.0.

### Statistical Analysis

Significant differences in Kaplan-Meier survival curves were calculated using the log rank test*. In vitro* growth/decay rates were analyzed by nonlinear regression of log_10_ values, and linear slopes were compared by analysis of covariance (ANCOVA). The remainder of statistical analyses were performed on linear values. Unpaired t-tests were done to analyze *in vitro* growth dynamics in ThyB and lung cultures. Analyses of variance (ANOVA) were performed using a Dunnett correction for multiple comparisons (to WT D39) to analyze *in vivo* differences including, lung and blood bacterial loads, viral loads, immune cells, cytokines, and chemokines (GraphPad Prism 7.0c). The confidence interval of significance was set to 95%, and p values less than 0.05 were considered significant.

## Contributions

AMS, JAM, and JR conceived the experiments. AMS, AI, CB, and MJ generated the mutant libraries. DR, RC, and TvO completed the sequence analyses. APS and LL generated SGD mutant bacteria. AMS, APS, LL, and MJ performed *in vitro/in vivo* experiments. AMS, APS, LL, and GH performed the data analysis. PV performed the histological analysis. AMS and APS wrote the manuscript.

## Acknowledgements

This work was supported by NIH grant AI100946, AI125324, and AI139088 (AMS), PATRIC DBP (AMS, JR, JAM), ALSAC (APS, LCL, SW, RC, AI, CB, PV, DR, MDLJ, JAM, JR), NIH grant R25CA23944 (GH), and NIH grants R01AI110724, U01AI124302, and R21AI117247 (TvO).

