## Supplementary Information for "Dynamic pneumococcal genetic adaptations support bacterial growth and inflammation during coinfection with influenza"

### Tn-seq Sample Preparation and Illumina Sequencing

Genomic DNA from the frozen pellets from each of the three time points (pre-selection (inoculum) ($t_1$) and post-selection (after infection) at 12 h ($t_2$) or 24 h ($t_3$) pbi) was digested overnight at 37°C with MmeI (NEB), the 5' phosphate group was removed with Calf intestinal alkaline phosphatase (NEB) after which DNA was extracted with the Geneaid Small Fragment DNA kit and dissolved in $H_2O$. An adapter was ligated with T4 DNA ligase (NEB) onto the overhang left by MmeI after which a PCR was performed with the adapter-ligated samples as template. One primer was complementary to the mini-transposon inverted repeat sequence and one primer was to the adapter (Table S1). The resulting PCR product was 140 bp in length and was amplified with the following parameters: 95°C for 30 sec, 26 cycles of 10 sec at 95°C, 25 sec at 55°C and 45 sec at 72°C, 1 cycle of 10 min at 72°C, and held at 4°C. The PCR product was purified using the Agencourt AMPure XP kit, dissolved in $H_2O$ and sequenced in rapid run mode on an Illumina HiSeq 2000 according to the manufacturers protocol (Illumina). A 6-nucleotide barcode sequence was included with the adapter so that harvested libraries coud be multiplexed. Following 30 sequencing cycles, raw data is extracted, split into different samples based on the 6-nucleotide-barcode sequences and stripped from the barcode and four nucleotides of the adapter sequence. This resulted in 5-15x$10^6$ pneumococcal specific reads per flow cell lane.

#### Fitness Calculations

Fitness calculations were performed as previously described (1, 2). Following sequencing, reads were mapped to the D39 genome using Bowtie (3). Bowtie parameters (-$m_1$–$n_1$–best) were set so that reads could contain a single mismatch but were only allowed if they mapped to a unique location. If mapping to multiple sites was possible, the read was excluded from the analyses. Approximately 8% of the reads had to be discarded because they could be mapped to multiple sites such as endogenous transposon related genes or other repeated sequences (6%) or could not be mapped to anywhere and were categorized as junk sequences (2%). Insertions that mapped to a location within the first 5% or the last 10% of a gene were removed from the analysis to minimize the influence of truncated functional genes. On average, 250 reads were mapped per insertion/time point. Only insertions with >15 reads in the inoculum were included in the analyses because insertions with a low number of reads slightly fluctuate over time and can influence the data disproportionately. The data were normalized to the total number of sequenced reads per time point (normalization factors were between 0.92 and 1.06). Fitness was calculated as the change in the number of reads at a specific location over time (see Main Text). Following fitness calculations, the values were normalized against a set of 'neutral' genes. These genes have no fitness effect and consist of pseudo genes and degenerate transposon related sequences. The same factor was then used to normalize the remaining dataset and make all fitness values relative to the WT D39 background. The normalization factors used for all datasets were small and were between 0.98 and 1.09. Each insertion was used to calculate the average fitness and standard deviation of the gene. A weighted average was used to control for fitness deviations due to insertions with small numbers of reads (<50 reads). This resulted in a small increase in replicate correlation and lower standard deviation.

To determine *in vivo* fitness and account for random loss of mutants during inoculation, the same proportion of insertion mutants that disappeared during *in vivo* selection were removed from the total number of insertions for each gene. The resulting set of insertions was then reanalyzed and the fitness recalculated. The resulting fitness ($W_i$, see Main Text) for each gene represents the growth rate per generation, which enables direct comparisons between experiments. To determine which genes differed with statistical significance, the fitness in influenza-infected mice was compared to the fitness in PBS-infected mice. Statistical significance was established when (i) fitness was composed of at least four data points, (ii) fitness deviated by at least 20%, and (iii) a one sample *t*-test with Bonferroni correction had a p-value less than 0.05.

**Table S1: Primers and Amplification Cycles:** Primers and amplification cycle used to (A) sequence the 140 bp Tn-seq region, (B) generate regions flanking the D39 locus targeted for deletion with overhangs complementary to ERM resistance cassette (Regions A and B) where the sequences in black are complementary to D39 genomic DNA and the sequences in red are complementary to ERM resistance cassette DNA, (C) delete the target locus by SOEing PCR to generate SGD mutants (D39Δ*cbiO1*, D39Δ*purD*, D39Δ*1414*, D39Δ*1098*, and D39Δ*proB*), and (D) confirm ERM resistance cassette insertion and target locus deletion.

| A) Tn-Seq | | |
|---|---|---|
| Primers | Sequence (5'-3') | Amplification Cycle |
| P1-M6-MmeI | CAAGCAGAAGACGGCATACGAAGACCGGGGACTTATCATCCAACCTGT | 95°C (30sec),<br>[95°C (10sec), 55°C (25sec), 72°C (45sec) x25], 72°C (10min),<br>4°C (hold) |
| ADPT-Tnseq-PCR | AATGATACGGCGACCACCGAGATCTACACTCTTTCCCTACACGACGCTCTTCCGATCT | |

| B) SOEing Regions | | | | |
|---|---|---|---|---|
| Locus | Product | | Sequence (5'-3') | Amplification Cycle |
| **SPD0058** 1.26kb: base 55644-56906 | Region A | F | CGCTTTGATTGCAGAATACTTCACAGC | 95°C (2min), [95°C (40sec), 48°C (30sec), 72°C (80sec) x40], 72°C (7min),<br>4°C (hold) |
| | | R | GTTTGCTTCTAAGTCTTATTTCCTCCTCAACCTCTTTTGCGAATTATTTACC | |
| | Region B | F | GAGTCGCTTTTGTAAATTTGGAGATATAAGAATAACGCGCCGTAGTCGC | |
| | | R | CGACATTTTCAAACTCGTAAGTGAGG | |
| **SPD1098** 2.17kb: base 1127189-1129354 | Region A | F | GCCAAAAGAACCACCTGATAGCC | 95°C (2min), [95°C (40sec), 48°C (30sec), 72°C (90sec) x40], 72°C (7min),<br>4°C (hold) |
| | | R | GTTTGCTTCTAAGTCTTATTTCCTTTTAGTCTCCTTTTCCGAATATTCTC | |
| | Region B | F | GAGTCGCTTTTGTAAATTTGG TGGCAAAATTAAAAATTGATGTAAATGATTTAC | |
| | | R | GCCAAATCAAGGATGGACTCG | |
| **SPD1414** 1.23kb: base 1433092-1434318 | Region A | F | CCAACTGCAAAGACACCAGGAACG | 95°C (2min), [95°C (40sec), 48°C (30sec), 72°C (90sec) x40], 72°C (7min),<br>4°C (hold) |
| | | R | GTTTGCTTCTAAGTCTTATTTCCAAAACCTCCTATTTTCGAACATTTATTC | |
| | Region B | F | GAGTCGCTTTTGTAAATTTGGTTTTCTGTAGAAATGGGGCTATCTTTTGC | |
| | | R | CCGATACGGTGAACATAACTCTCAGG | |

| | | | | |
|---|---|---|---|---|
| **SPD2047** 840bp: base 2023773-2024612 | Region A | F | GCTATTATTTTCTTGCATATTACATTGG | 95°C (2min), [95°C (40sec), 48°C (30sec), 72°C (90sec) x40], 72°C (7min), 4°C (hold) |
| | | R | GTTTGCTTCTAAGTCTTATTTCCAGCTTATCCTCTAGCTCACTTTCTGTC | |
| | Region B | F | GAGTCGCTTTTGTAAATTTGGTATGATTTTGGGGCGTTATATCCCAGGG | |
| | | R | CTCTATCAACCAATGAGATTCCATCTCC | |
| **SPD0822** 1.1kb: base 840495-841604 | Region A | F | GCTGGTTGGCATAAGAGTGC | 95°C (2min), [95°C (40sec), 48°C (30sec), 72°C (120sec) x40], 72°C (7min), 4°C (hold) |
| | | R | GTTTGCTTCTAAGTCTTATTTCCAGTCTCACTCCGATTATTCATATTTATCC | |
| | Region B | F | GAGTCGCTTTTGTAAATTTGGAGGTAAACTATGGTGAGTAGAC | |
| | | R | GCTGTGGGATTCAATGTGC | |
| **ERM** | | F | GGAAATAAGACTTAGAAGCAAAC | 95°C (2min), [95°C (40sec), 48°C (30sec), 72°C (90sec) x40], 72°C (7min), 4°C (hold) |
| | | R | CCAAATTTACAAAAGCGACTC | |
| **C) SOEing PCR** | | | | |
| Locus | | | Sequence (5'-3') | Amplification Cycle |
| **SPD0058** | | F | CGCTTTGATTGCAGAATACTTCACAGC | [94°C (30sec), 50°C (30sec), 72°C (3.25min) x34], 72°C (5min), 4°C (hold) |
| | | R | CGACATTTTCAAACTCGTAAGTGAGG | |
| **SPD1098** | | F | GCCAAAAGAACCACCTGATAGCC | [94°C (30sec), 50°C (30sec), 72°C (3.50min) x34], 72°C (10min), 4°C (hold) |
| | | R | GCCAAATCAAGGATGGACTCG | |
| **SPD1414** | | F | CCAzACTGCAAAGACACCAGGAACG | [94°C (30sec), 50°C (30sec), 72°C (3.50min) x34], 72°C (10min), 4°C (hold) |
| | | R | CCGATACGGTGAACATAACTCTCAGG | |

| | | | |
|---|---|---|---|
| **SPD2047** | F | GCTATTATTTTCTTGCATATTACATTGG | [94°C (30sec), 48°C (30sec), 72°C (3.50min) x34], 72°C (5min), 4°C (hold) |
| | R | CTCTATCAACCAATGAGATTCCATCTCC | |
| **SPD0822** | F | GCTGGTTGGCATAAGAGTGC | [94°C (30sec), 50°C (30sec), 72°C (4min) x34], 72°C (10min), 4°C (hold) |
| | R | GCTGTGGGATTCAATGTGC | |
| **D) Confirmation of Locus Deletion** | | | |
| Locus | | Sequence (5’-3’) | Amplification Cycle |
| **SPD0058** | F | GCCCTTGCTGCTGGTATCGTGG | 95°C (2min), [95°C (40sec), 48°C (30sec), 72°C (60sec) x40], 72°C (7min), 4°C (hold) |
| | R | GGCTTGACAATGGTGTCAACCG | |
| **SPD1098** | F | GCTGTAGGAAGTTTTGCTTTCGG | 95°C (2min), [95°C (40sec), 48°C (30sec), 72°C (60sec) x40], 72°C (7min), 4°C (hold) |
| | R | CCGATTGGAGTTCCAGAGATTGG | |
| **SPD1414** | F | GCAATCTTTTGTTTGGGCTTATCGG | 95°C (2min), [95°C (40sec), 48°C (30sec), 72°C (60sec) x40], 72°C (7min), 4°C (hold) |
| | R | GCTGGAATCAAAGAAAAACCAGC | |
| **SPD2047** | F | CGGATGTTTCTTTGACGATTGAAGATGGC | 95°C (2min), [95°C (40sec), 48°C (30sec), 72°C (60sec) x40], 72°C (7min), 4°C (hold) |
| | R | CCCTAGAGGATCTAGACCAGCTGTTGGC | |
| **SPD0822** | F | CGTCAAATCGTTTCTGC | 95°C (2min), [95°C (40sec), 48°C (30sec), 72°C (60sec) x40], 72°C (7min), 4°C (hold) |
| | R | GCCATTGTTTCTGGGTACG | |

### Pathogenicity of Single-gene Deletion Bacterial Mutants

Table S2 summarizes significant differences in Kaplan-Meier survival curves, the $\log_{10}$ changes, and statistical comparisons in lung bacteria, bacteremia, and viral load during naïve infection and IAV-coinfection with each of the SGD mutant bacteria, compared to WT D39.

**Table S2: Summary of Pathogenicity of SGD Mutant Bacteria**. The $\log_{10}$ change ($\Delta$) in the levels of lung bacteria, blood bacteria, and lung virus from mice (groups of 5) either mock-infected (PBS) or IAV-infected (75 $TCID_{50}$ PR8) followed 7 d later by $10^6$ CFU of the indicated bacteria. Comparisons were made between average $\log_{10}$ bacteria or virus at 4 h pbi and 24 h pbi. Significance of the analysis of variance (ANOVA) using a Dunnett correction for multiple comparisons (to WT D39) is indicated in bold. The percent increase in survival of animals (groups of 10) either mock-infected (PBS) or IAV-infected (75 $TCID_{50}$ PR8) followed 7 d later by $10^6$ CFU of the indicated bacteria. Significant differences in Kaplan-Meier survival curves by the log rank test are indicated in bold. Significance is *p<0.05, **p<0.01, ***p<0.005, ****p<0.0001.

| **Bacteria** | **$\Delta$ Lung Bacteria ($\log_{10}$)** | | | | **$\Delta$ Blood Bacteria ($\log_{10}$)** | | | | **$\Delta$ Virus ($\log_{10}$)** | | **Survival** | |
|---|---|---|---|---|---|---|---|---|---|---|---|---|
| | **PBS + Bacteria** | | **PR8 + Bacteria** | | **PBS + Bacteria** | | **PR8 + Bacteria** | | **PR8 + Bacteria** | | **PBS** | **PR8** |
| | **4 h** | **24 h** | **4 h** | **24 h** | **4 h** | **24 h** | **4 h** | **24 h** | **4 h** | **24 h** | | |
| **D39Δ1414** | **-0.61******** | -2.71 | **-0.94******** | **-0.83**** | -3.79 | -5.87 | -1.65 | **-4.97**** | 0.02 | 0.43 | **100%******** | **80%******** |
| **D39ΔcbiO 1** | **-1.36******** | -5.86 | **-2.05******** | **-2.11**** | -2.31 | -6.56 | -1.38 | **-2.35**** | 0.15 | 0.29 | **90%******** | **40%****** |
| **D39ΔpurD** | **-0.74******** | -0.47 | **-0.44******** | **-0.87*** | -1.49 | -1.28 | -0.40 | **-2.64**** | -0.83 | 0.26 | **100%******** | **60%****** |
| **D39Δ1098** | **-0.59******** | -2.17 | **-0.76******** | **-0.78**** | -3.78 | -3.51 | -1.69 | **-2.25**** | -0.83 | -0.93 | **100%******** | **90%******** |
| **D39ΔproB** | **-1.07******** | -1.70 | **-1.08******** | **-0.86*** | -3.21 | -6.56 | -1.86 | **-2.98**** | -0.04 | -0.19 | **40%******** | **0%[†]******** |

† Mean survival time at 7 d pbi

### *In vitro* Growth of Single-gene Deletion Bacterial Mutants following Metabolic Starvation

**Error! Reference source not found.** shows the growth of each SGD mutant bacteria and WT D39 in cultures supplemented with lung homogenate supernatants (s/n) from mock- or IAV-infected mice, following 5 h of metabolic starvation or from culture initiation.

### Kinetics of Pulmonary Cytokines, Chemokines, and Immune Cells during Naïve infection and IAV-Coinfection with Single-gene Deletion Bacterial Mutants

The absolute $\log_{10}$ picograms (pg) of all measured cytokines and chemokines in mock- and IAV-infected animals are in Fig S2-S4. A heat map of the fold change in cytokines and chemokines is in Fig 5 of the main text. Pulmonary immune cells, quantified by flow cytometry according to the gating scheme in Fig S5, are shown in Fig S6. Additional cell kinetics are shown in Fig 6 of the main text.

### Quantification of Pneumococcus and Neutrophils by IHC Staining

Figure S7 shows the percent of the lung positive for pneumococcus or neutrophil antigen in serial sections of lungs at 24 h pbi from mice IAV-infected (75 $TCID_{50}$ PR8) followed 7 d later with $10^6$ CFU of wild-type or SGD bacteria.

### Bacterial Loads and Survival Following Varied Doses of Wild-type Bacteria

Figure S8 shows bacterial loads and Kaplan-meier survival curves for WT D39 infections of mock- and IAV-infected animals. Significantly reduced bacterial loads at 24 h pbi result in a 20% reduction in lethality in IAV-infected animals.

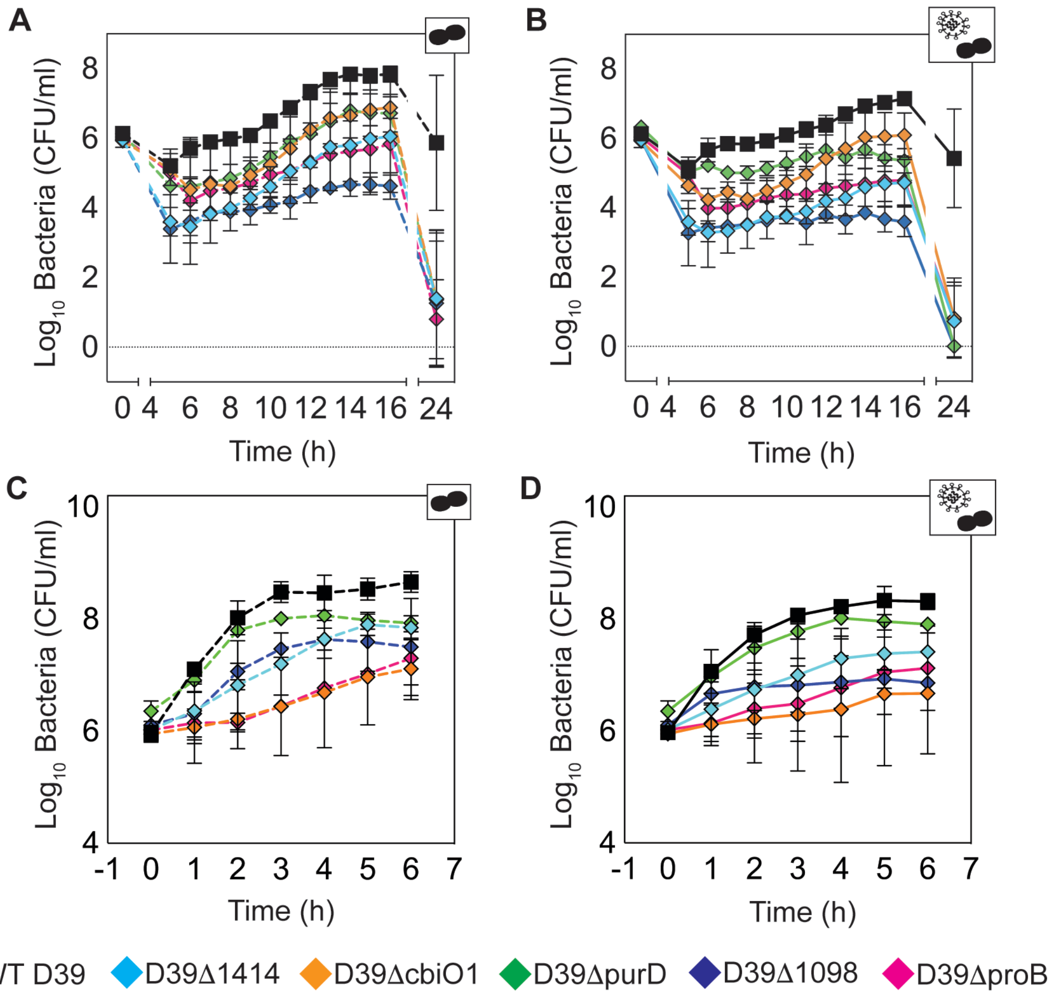


**Fig S1: *In vitro* Growth in Lung Supernatants.** Bacteria were grown at 37°C in either 1 ml of PBS for 5 hours and then supplemented with 0.5 ml lung homogenate s/n (Panels A and B), or cultured continuously in lung homogenate s/n (Panels C and D). Lung homogenates were collected 7 d pi from mice mock-infected (PBS) (Panels A and C) or IAV-infected (75 $TCID_{50}$ PR8) (Panels B and D) . Cultures were sampled hourly (50 µl) and samples were serially diluted in PBS and plated on TSA (WT) (squares) or TSA-ERM (diamonds) plates. Bacterial titers were normalized to the total culture volume. From hours 7-16 of culture, each SGD mutant was significantly lower than WT D39 ($p<0.05$) after the addition of lung s/n, except for D39Δ*purD* at 11 hours in mock-infected lung s/n

(Panel A) and D39Δ*cbiO1*, D39Δ*purD*, and D39Δ*proB* at 12 h in IAV-infected lung s/n (Panel B). Bacterial titers of SGD mutants were not significantly lower than WT D39 after 6 h of continuous culture in lung s/n from mock-infected (Panel C) or IAV-infected (Panel D) mice.

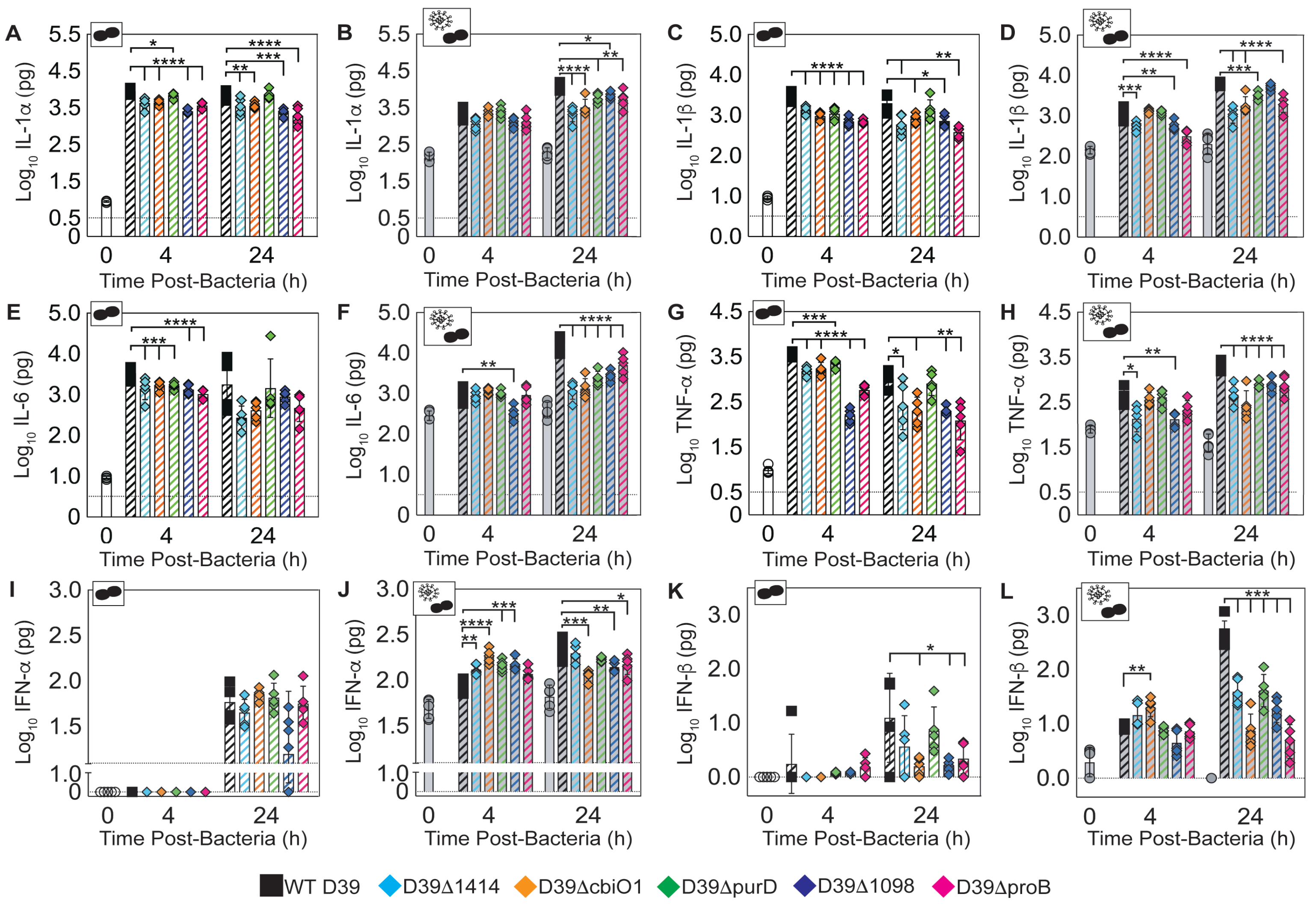


**Fig S2: Pulmonary Cytokine Kinetics.** Kinetics at 4 h and 24 h pbi of IL-1α (A-B), IL-1β (C-D), IL-6 (E-F), TNF-α (G-H), IFN-α (I-J), and IFN-β (K-L) from mice either mock-infected (PBS) (Panels A, C, E, G, I, K) or IAV-infected (75 $TCID_{50}$ PR8)

(Panels B, D, F, H, J, L) followed 7 d later with $10^6$ CFU of the indicated bacteria. Each symbol (circles, squares, or diamonds) represents a single mouse, and the bars are the geometric mean ± standard deviation (SD) from 5 mice/group. Mice were either uninfected (open white), IAV-infected only for 7 d (T=0) or 8 d (T=24) (solid grey), bacteria-infected (open hashed, colored), or IAV-bacteria coinfected (solid hashed, colored). Cartoons indicating infection status of study group (bacteria alone or virus plus bacteria) are in the upper right corner of each graph. Dashed line indicates the lower limit of detection (LOD), undetectable amounts are plotted at 0 $\log_{10}$ pg, and values extrapolated by the 4PL analysis appear between 0 $\log_{10}$ pg and the LOD. Signficance is indicated as *p<0.05, **p<0.01, ***p<0.005, ****p<0.0001. Additional cytokines are shown in Fig S3 and a heat map of cytokines and chemokines is in Fig 5 (Main Text).

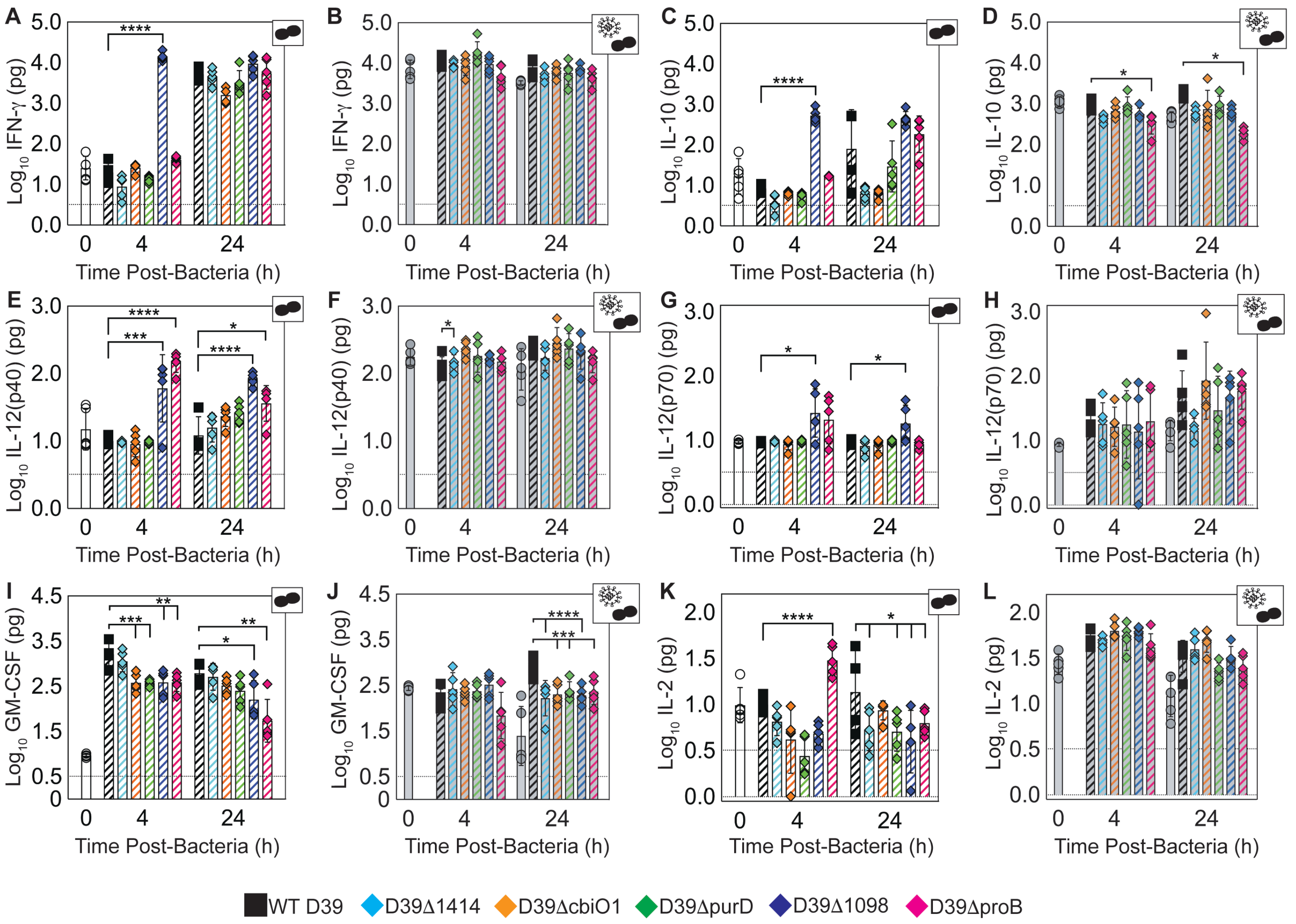


**Fig S3: Additional Cytokine Kinetics in the Lung.** Kinetics at 4 h and 24 h pbi of IFN-γ (A-B), IL-10 (C-D), IL-12(p40) (E-F), IL-12(p70) (G-H), GM-CSF (I-J), and IL-2 (K- L) from mice either mock-infected (PBS) (Panels A, C, E, G, I, K) or IAV-infected (75 $TCID_{50}$ PR8) (Panels B, D, F, H, J, L) followed 7 d later with $10^6$ CFU of the indicated bacteria. Each symbol (circles, squares, or diamonds) represents a single mouse, and the bars are the geometric mean ± standard deviation (SD) from 5 mice/group. Mice were either uninfected (open white), IAV-infected only for 7 d (T=0) or 8 d (T=24), bacteria-infected (open hashed, colored), or IAV-bacteria coinfected (solid hashed, colored). Cartoons indicating infection status of study group (bacteria alone or virus plus bacteria) are in the upper right corner of each graph. Dashed line indicates the lower limit of detection (LOD), undetectable amounts are plotted at 0 $\log_{10}$ pg, and values extrapolated by the 4PL analysis appear between 0 $\log_{10}$ pg and the LOD. Signficance is indicated as *p<0.05, **p<0.01, ***p<0.005, ****p<0.0001. Additional cytokines are shown in Fig S2 and a heat map of cytokines and chemokines is in Fig 5 (Main Text).

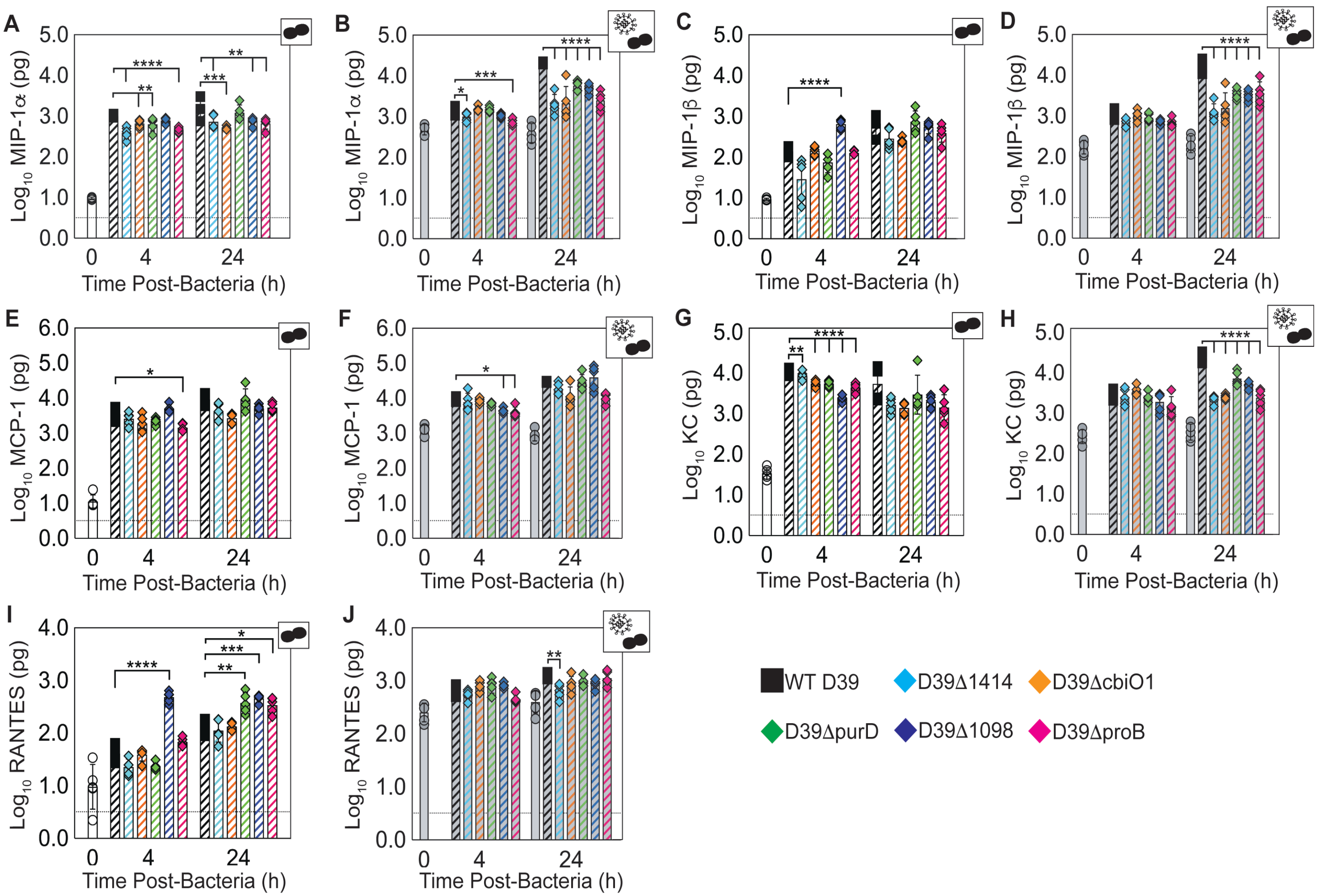


**Fig S4: Pulmonary Chemokine Kinetics.** Kinetics at 4 h and 24 h pbi of MIP-1α (A-B), MIP-1β (C-D), MCP-1 (E-F), KC (G-H), and RANTES (I-J) from mice either mock-infected (PBS) (Panels A, C, E, G, I) or IAV-infected (75 $TCID_{50}$ PR8)

(Panels B, D, F, H, J) followed 7 d later with $10^6$ CFU of the indicated bacteria. Each symbol (circles, squares, or diamonds) represents a single mouse, and the bars are the geometric mean ± standard deviation (SD) from 5 mice/group. Mice were either uninfected (open white), IAV-infected only for 7 d (T=0) or 8 d (T=24) (solid grey), bacteria-infected (open hashed, colored), or IAV-bacteria coinfected (solid hashed, colored). Cartoons indicating infection status of study group (bacteria alone or virus plus bacteria) are in the upper right corner of each graph. The dashed line indicates the lower limit of detection (LOD). Signficance is indicated as *$p<0.05$, **$p<0.01$, ***$p<0.005$, ****$p<0.0001$. A heat map of cytokines and chemokines is in Fig 5 (Main Text).

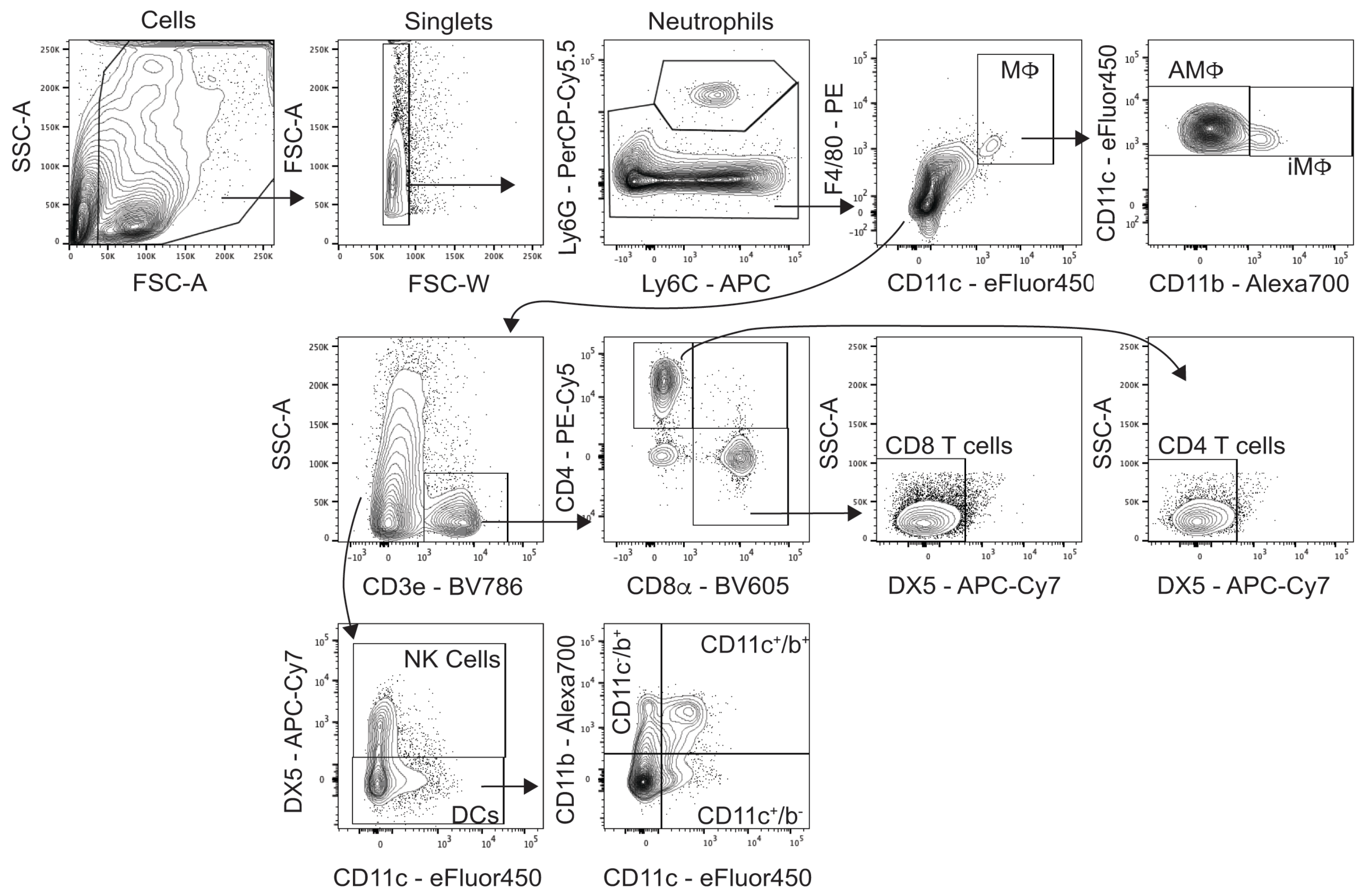


**Fig S5: Flow Cytometry Gating Scheme for Lung Cell Analysis.** Live cells were first gated on forward scatter (FSC-A) and side scatter (SSC-A) then as singlets. Neutrophils (Ly6G$^{hi}$) were then gated and excluded from remaining parent populations. Macrophages (MΦ) were gated as CD11c$^{hi}$F4/80$^{hi}$ with alveolar macrophages (AMΦ) sub-gated as CD11b$^{-}$

MHC-II$^{low}$ and inflammatory macrophages (iM$\Phi$) as CD11b$^{+}$MHC-II$^{mid/hi}$. Following M$\Phi$ exclusion, T cells were gated as CD3e$^{+}$ and subgated into CD8$^{+}$ T cells (CD3$^{+}$CD8$^{+}$CD4$^{-}$DX5$^{-}$) and CD4$^{+}$ T cells (CD3$^{+}$CD8$^{-}$CD4$^{+}$DX5$^{-}$) populations. Of the cells that were CD3e$^{-}$, natural killer (NK) cells were gated as CD3$^{-}$DX5$^{+}$ and dendritic cells (DC) as CD3$^{-}$DX5$^{-}$. DCs were subgated as CD11c$^{+}$CD11b$^{-}$ DCs, CD11c$^{+}$CD11b$^{+}$ DCs, and CD11c$^{-}$CD11b$^{+}$ DCs.

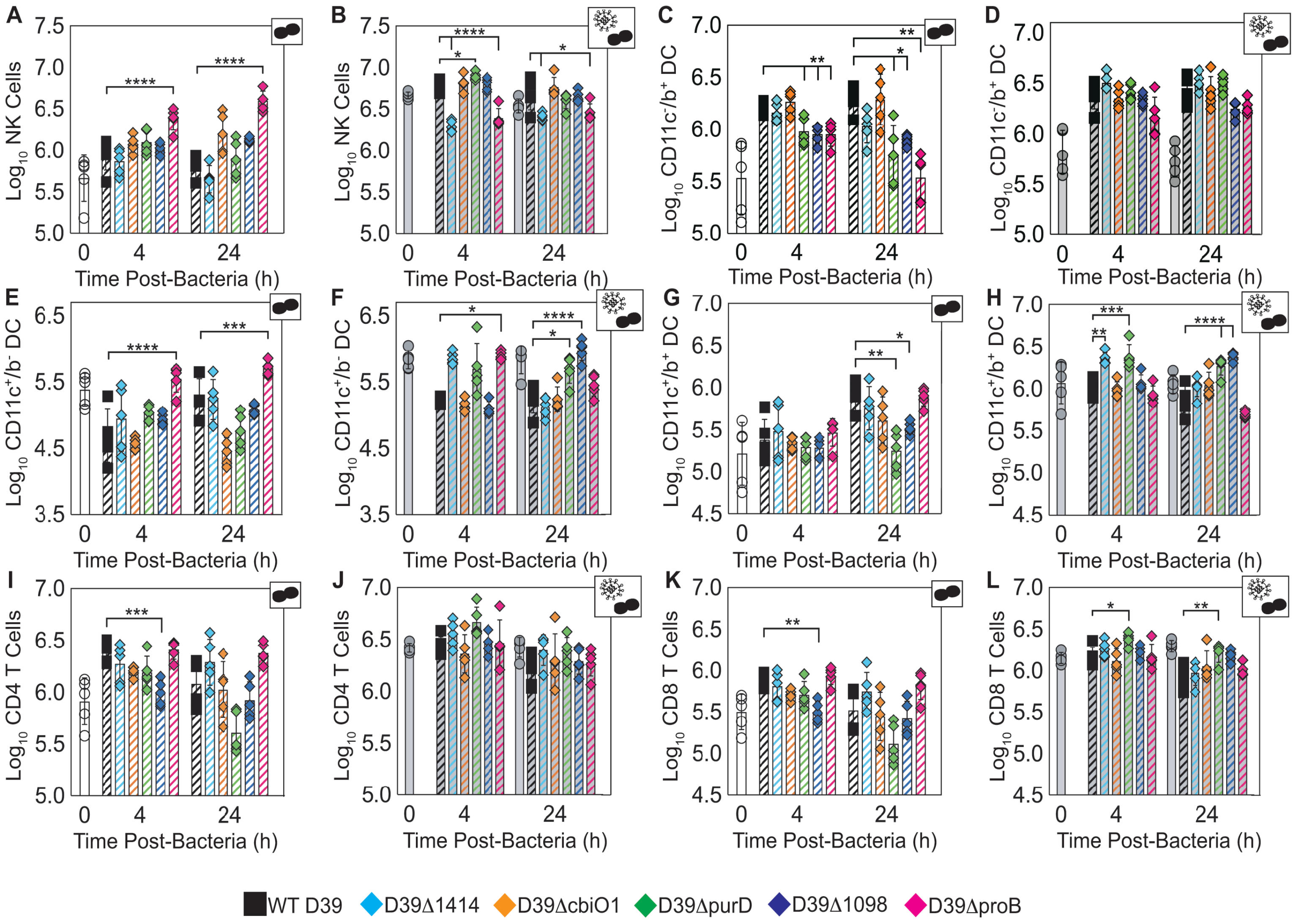


**Fig S6: Additional Pulmonary Immune Cell Kinetics**. Kinetics at 4 h and 24 h pbi of NK cells (A-B), DCs (C-H), and T cells (I-L) from mice either mock-infected (PBS) (Panels A, C, E, G, I, K) or IAV-infected (75 $TCID_{50}$ PR8) (Panels B, D, F,

H, J, L) followed 7 d later with $10^6$ CFU of the indicated bacteria. Each symbol (circles, squares, or diamonds) represents a single mouse, and the bars are the geometric mean ± standard deviation (SD) from 5 mice/group. Mice were either uninfected (open white), IAV-infected only for 7 d (T=0) or 8 d (T=24) (solid grey), bacteria-infected (open hashed, colored), or IAV-bacteria coinfected (solid hashed, colored). Cartoons indicating infection status of study group (bacteria alone or virus plus bacteria) are in the upper right corner of each graph. Signficance is indicated as *p<0.05, **p<0.01, ***p<0.005, ****p<0.0001. Additional cells are shown in Fig 6 (Main Text) and the flow cytometry gating scheme is in Fig S5.

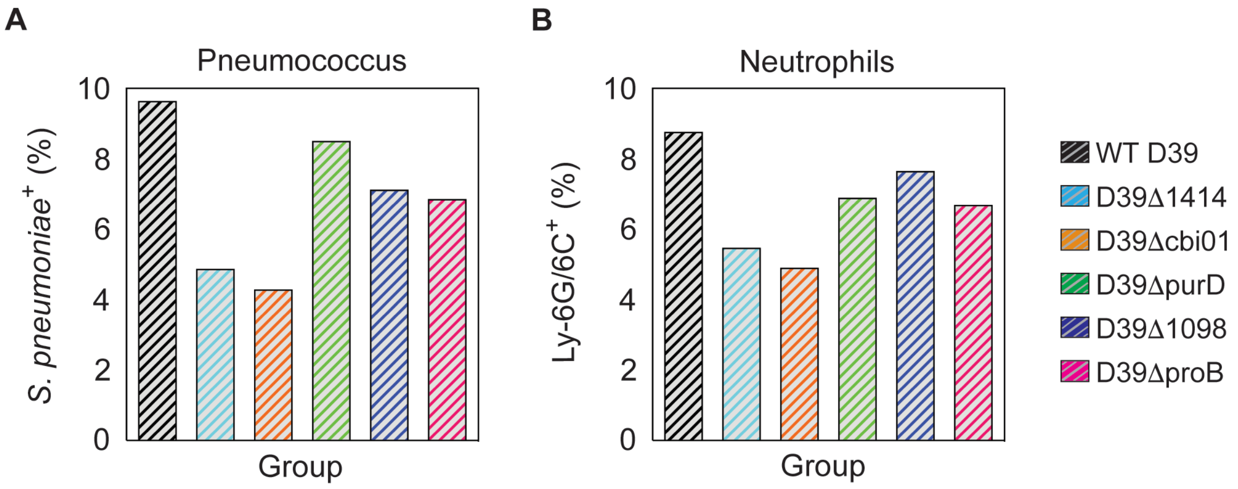


**Figure S7: Quantification of IHC Staining During IAV-Pneumococcal Coinfection.** Serial sections of lungs at 24 h pbi from mice IAV-infected (75 $TCID_{50}$ PR8) followed 7 d later with $10^6$ CFU of the indicated bacteria were stained for pneumococcus or neutrophil Ly6G/6C antigen. The immunohistochemistry recipe #L006-02 provided in MIPAR v3.0 was used to quantify staining in the entire lung section that representative images in Figure 7C (Main Text) were taken from, and are presented as percent area positive for pneumococcal (Panel A) and Ly6G/6C (Panel B) antigen.

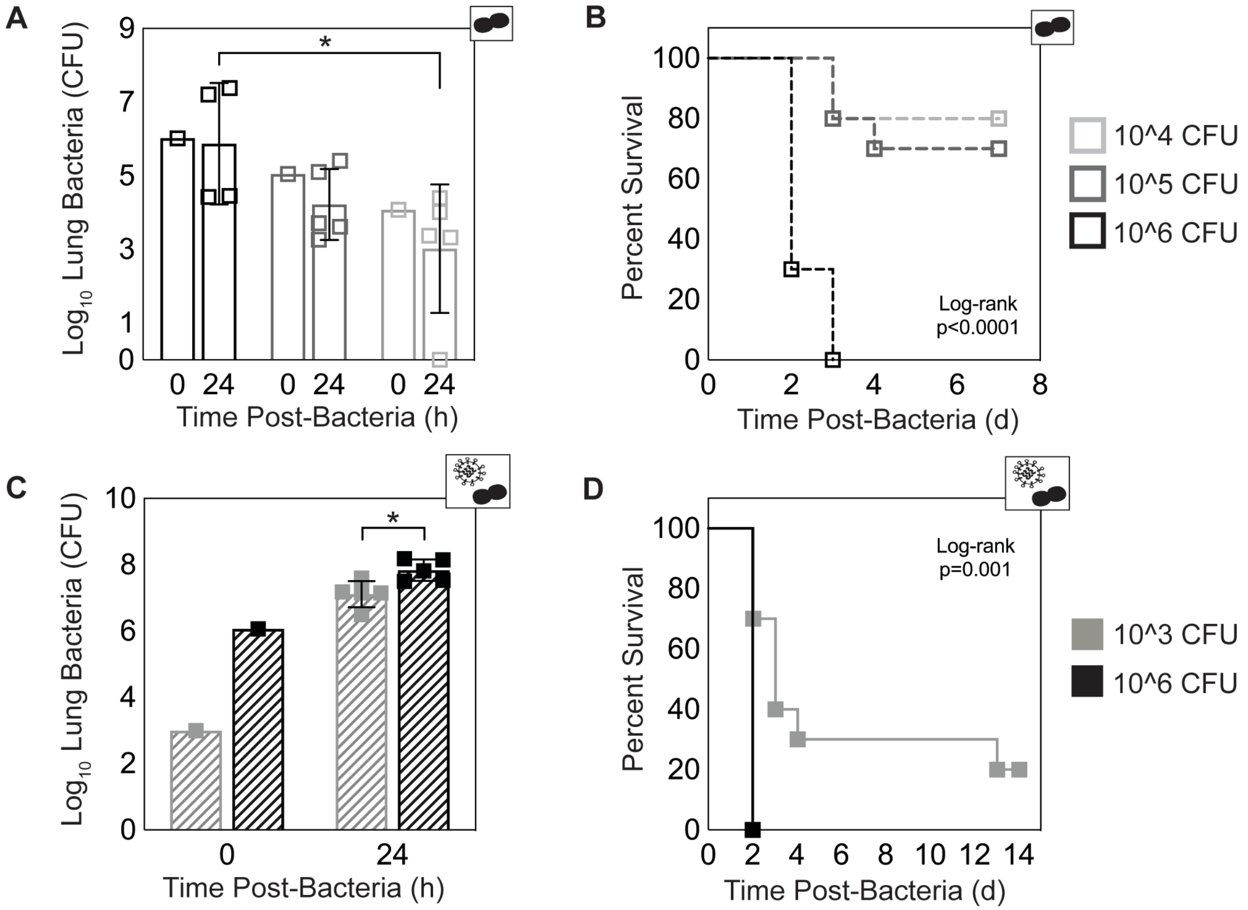


**Fig S8: Lung Bacterial Titers and Pathogenicity of Infection with Varied Doses of WT D39.** Lung bacterial titers (Panels A, C) and Kaplan-Meier survival curves (B-D) of mice mock-infected (PBS) (Panels A-B) or IAV-infected (75 $TCID_{50}$ PR8) (Panels C-D) followed 7 d later with the indicated dose of WT D39 bacteria. Survival curves are significantly different for each dose compared to $10^6$ CFU. Signficant differences in bacterial titers at 24 h pbi are indicated as *p<0.05. Each symbol represents a single mouse, and the bars are the geometric mean ± standard deviation (SD) from 5 mice/group. Cartoons indicate infection status of study group (bacteria alone or virus plus bacteria).
